# A model of resource partitioning between foraging bees based on positive and negative associations

**DOI:** 10.1101/2020.11.13.381012

**Authors:** Thibault Dubois, Cristian Pasquaretta, Andrew B. Barron, Jacques Gautrais, Mathieu Lihoreau

## Abstract

Central place foraging pollinators tend to develop multi-destination routes (traplines) to visit patchily distributed plant resources. While the formation of traplines by individual pollinators has been studied in details, how populations of foragers exploit resources in a common area is an open question, difficult to address experimentally. We explored conditions for the emergence of resource partitioning among traplining bees using agent-based models built from experimental data of bumblebees foraging on artificial flowers. In the models, bees learn to develop routes as a consequence of feedback loops that change their probabilities of moving between flowers. While a positive reinforcement of movements leading to rewarding flowers is sufficient for the emergence of resource partitioning when flowers are evenly distributed, addition of negative reinforcement of movements leading to unrewarding flowers is necessary when flowers are patchily distributed. In environments with more complex spatial structure, the negative experiences of individual bees on flowers favour spatial segregation and efficient collective foraging.

## Introduction

Foraging animals are expected to self-distribute on food resources in order to minimize competition and maximize their individual net energy gain (Fretwell and Lucas 1969; Giraldeau and Caraco 2000). Resource partitioning between individuals of different species is well documented, and often results from functional (Fründ *et al*. 2010; 2013) or behavioural (Nagamitsu and Inoue 1997; Valdovinos *et al*. 2016) specializations. By contrast, how individuals of the same species optimally interact to exploit resources in a common foraging area is less understood (Johst *et al*. 2008; Tinker *et al*. 2012).

For pollinators, such as bees that individually exploit patchily distributed floral resources in environments with high competition pressure, efficient resource partitioning appears a prodigious problem to solve, as it involves assessing the quality of food resources, their spatial distribution, their replenishment rate, and the activity of other pollinators. As central place foragers, bees often visit familiar feeding sites (plants or flower patches) in a stable sequence called “trapline” (Janzen 1971; Thomson *et al*. 1997). Individual bees with exclusive access to an array of artificial flowers tend to develop traplines minimizing travel distances to visit all the necessary flowers to fill their nectar crop and return to the nest (e.g. bumblebees: Ohashi *et al*. 2008, Lihoreau *et al*. 2012a, Woodgate *et al*. 2017; honey bees: Buatois and Lihoreau 2016). This routing behaviour involves spatial memories that can persist several days (Lihoreau *et al*. 2010) or weeks (Thomson 1996).

How bees partition resources, when several conspecifics exploit the same foraging area, is however an open question. Experimentally the problem is challenging to address as it requires monitoring the movements of numerous bees simultaneously over large spatial and temporal scales. In theory, bees should develop individualistic traplines that minimize travel distances and spatial overlap with other foragers, thereby improving their own foraging efficiency and minimizing the costs of competition (Ohashi and Thomson 2005; Lihoreau *et al*. 2016). Best available data supporting this hypothesis come from observations of small numbers of bumblebees foraging on potted plants (e.g. Makino and Sakai 2005; Makino 2013) or artificial flowers (e.g. Lihoreau *et al*. 2016; Pasquaretta *et al*. 2019) in large flight tents. In these experimental foraging conditions (with limited numbers of bees and feeding sites), foragers tend to avoid spatial overlaps as a consequence of competition by exploitation (when bees visited empty flowers) and interference (when bees interacted on flowers) (Pasquaretta *et al*. 2019).

Computational modelling is a powerful approach to further explore the mechanisms by which such partitioning might emerge from spatial learning and competitive interactions. At the individual level, trapline formation has been modelled using an iterative improvement algorithm where a bee compares the net length of the route it has just travelled (sum of the lengths of all movement vectors comprising the flower visitation sequence) to the length of the shortest route experienced so far (Lihoreau *et al*. 2012b). If the new route is shorter (or equivalent), the bee increases its probability of using all the movement vectors composing this route in its subsequent foraging bout. After several iterations, this route-based learning heuristic typically leads to the discovery and selection of a short (if not the shortest possible) trapline, thereby replicating observations in bees across a wide range of experimental conditions (Reynolds *et al*. 2013). Note however that this model makes the strong assumption that bees can compute, memorize and compare the lengths of multiple routes upon return to their nest. To address this issue, it was proposed that trapline formation could also emerge from vector-based learning (Le Moël *et al*. 2019), which is more parsimonious and plausible considering the current understanding of spatial computation in the insect brain (Stone *et al*. 2017). So far, however, none of these traplining algorithms have accounted for social interactions and current models of bee populations do not consider individual specificities of movements based on learning and memory (Becher *et al*. 2014; 2016; 2018). Thus presently, there has been no formal exploration of how resource partitioning between interacting bees might form.

Here, we investigated the behavioural mechanisms underlying resource partitioning among traplining bees by comparing predictions of agent-based models integrating route learning and social interactions. Recent work showed that resource partitioning in bats foraging on patchily distributed cacti can be explained by basic reinforcement rules, so that a bat that finds an abundant feeding site tends to return to this site more often than its conspecifics (Goldshtein et al. 2020). Since bees extensively rely on associative learnings to recognize flowers and develop foraging preferences (Giurfa 2013), we hypothesized that the combination of positive experiences (when a flower is full of nectar) and negative experiences (when a flower is unrewarding) could lead to the emergence of resource partitioning, when different bees learn to use spatially segregated routes (Lihoreau *et al*. 2016, Pasquaretta *et al*. 2019). First, we developed models implementing biologically plausible vector navigation based on positive and negative reinforcements of vectors leading to flowers and tested the independent and combined influences of these feedback loops on route learning by comparing simulations to published experimental data. Next, we explored how these simple learning rules at the individual level can promote complex patterns of resource partitioning at the collective level, using simulations with multiple foragers in environments with different resource distributions.

## RESULTS

### Overview of models

We designed models of agents (bees) foraging simultaneously in a common set of feeding sites (flowers) from a central location (colony nest) (see summary in Fig. 1). A full description of the models is available in the ODD protocol (Appendix S1). Briefly, each bee completes a succession of foraging trips (foraging bouts) defined as the set of movements and flower visits between the moment it leaves the nest until the moment it returns to it. Each bee initially moves between the different flowers using a distance-based probability matrix (Lihoreau *et al*. 2012b; Reynolds *et al*. 2013). The probability to move between each flower is then modulated each time the bee finds the flower rewarding (positive reinforcement) or unrewarding (negative reinforcement). Learning occurs after each flower visit (online learning). We implemented three models to explore different combinations of positive and negative reinforcements: model 1: positive reinforcement only (hereafter “pos.reinf”), model 2: negative reinforcement only (neg. reinf), model 3: positive and negative reinforcement (pos. + neg. reinf.). Model comparison thus informed about the effect of each of the rules separately and in combination.

**Figure 1.**
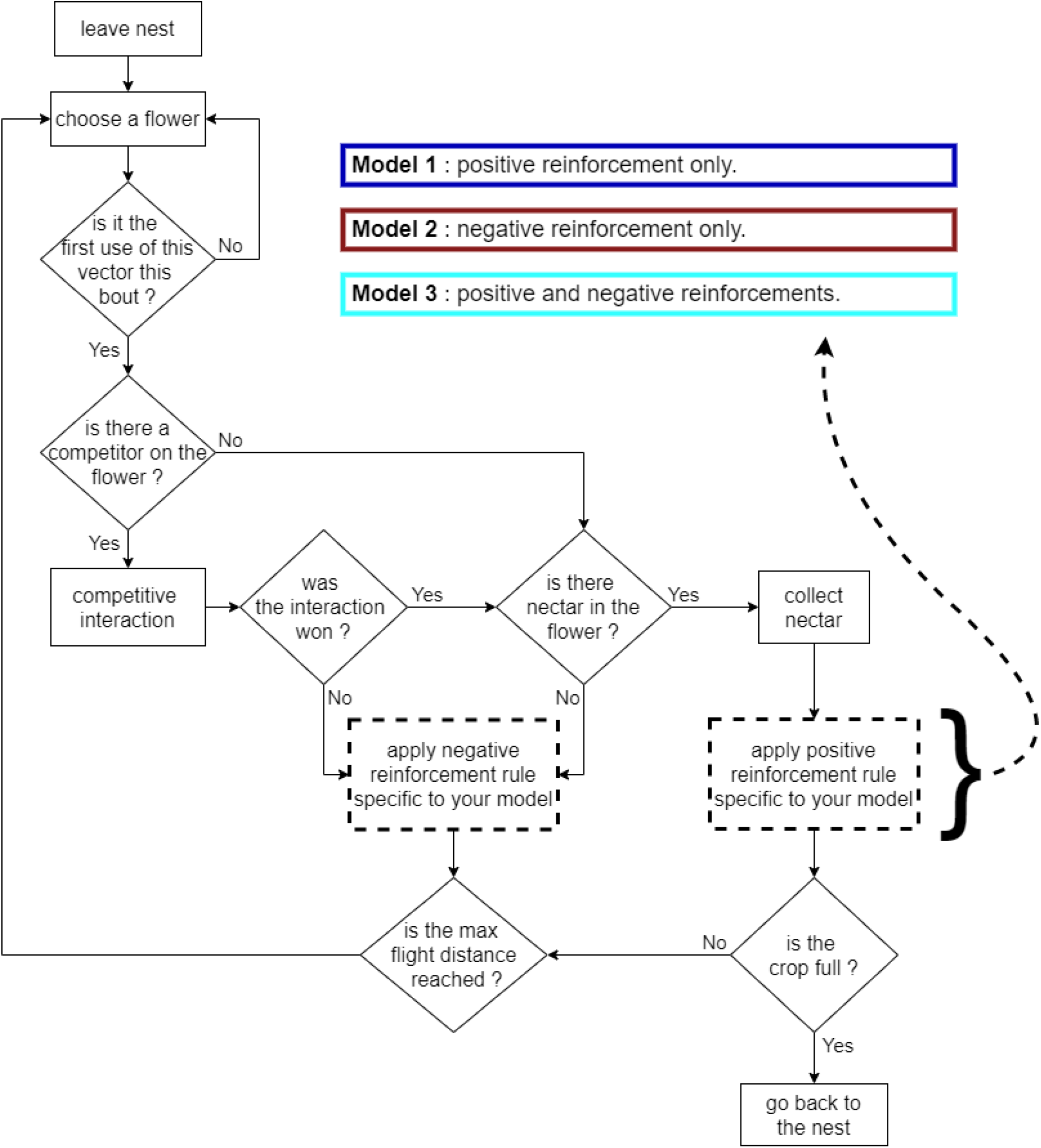
Flowchart summarizing the agent-based models. Rectangles represent actions performed by a bee. Diamonds indicate conditional statements. Arrows connect the different modules. The dashed rectangles are subject to the different rules of the three models.

### Simulations with one forager

We first tested the ability of our models to replicate trapline formation by real bees, by comparing simulations of a single forager to published experimental data in which individual bumblebees were observed developing traplines between five equally rewarding artificial flowers in a large open field (Lihoreau *et al*. 2012b; Woodgate *et al*. 2017). Lihoreau *et al*. (2012b) tested seven bumblebees in a regular pentagonal array (Fig. S1A), which we judged enough to run quantitative comparisons with model simulations. Woodgate *et al*. (2017) tested three bees in a narrow pentagonal array (Fig. S1B), which only enabled a qualitative comparison with the model simulations.

We assessed the ability of bees to develop efficient routes by computing an index of route quality (i.e. the squared volume of nectar gathered divided by the distance travelled; see Methods). For real bees, route quality increased significantly with time in the regular pentagonal array of flowers (Fig. 2A; GLMM_route quality_: *Est*. = 0.153 ± 0.023, *P* < 0.001). When comparing simulations to experimental data, there were no significant differences in trends with model 1 (Fig. 2A; pos. reinf.; GLMM_route quality_: *Est*. = −0.027 ± 0.023, *P* = 0.224) and model 3 (Fig. 2A; pos. + neg. reinf.; GLMM_route quality_: *Est*. = −0.022 ± 0.023, *P* = 0.339), meaning that simulated bees developed routes of similar qualities as real bees. However, route qualities of model 2 were significantly lower than the experimental data (Fig. 2A; neg. reinf. GLMM_route quality_: *Est*. = – 0.155 ± 0.023, *P* < 0.001). Similar trends were observed in the narrow pentagonal array of flowers (Fig. S5 and Appendix S3).

**Figure 2.**
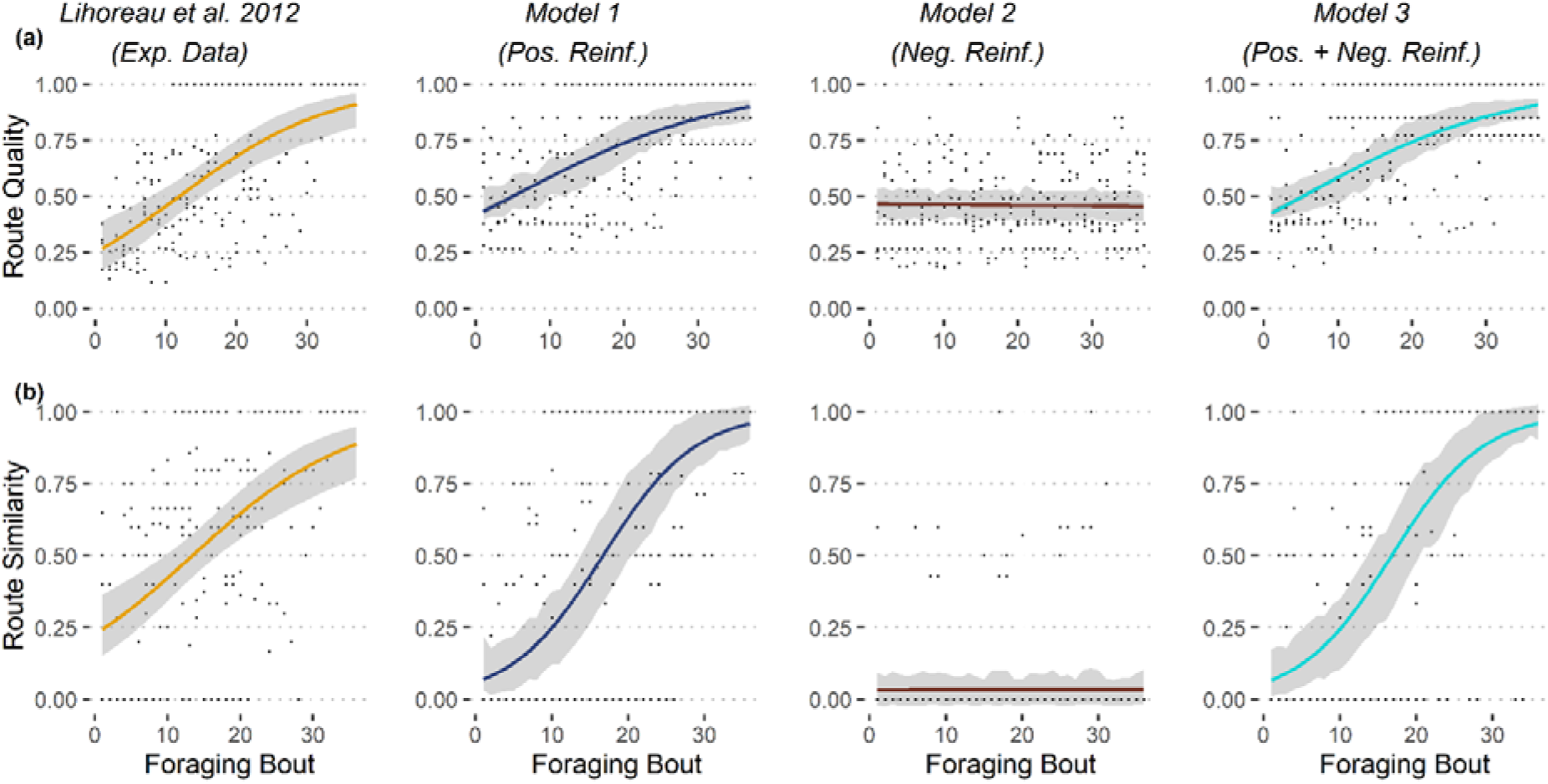
Comparisons of route qualities (a) and route similarities (b) between simulations and experimental data (regular pentagonal array of flowers as in Lihoreau *et al*. 2012b). See details of models in Fig. 1. For each dataset, we show the estimated average trends across foraging bouts (coloured curves), along with their 95% CI (grey areas) of the mean. For the sake of eye comparison, in the simulation plot we represent estimated 95% confidence intervals of the mean for a random subsample of 7 simulated bees (N = 7 bees in Lihoreau *et al*. 2012b). Average trends were estimated over 500 simulation runs, using GLMM Binomial model with bee identity as random effect (bee identity nested in simulation identity for simulated data).

We assessed the ability of bees to develop stable routes using an index of route similarity (i.e. computing the number and percentage of vectors shared between two successive routes; see Methods). Route similarity is set between 0 (the two routes are completely different) and 1 (the two routes are completely identical). For real bees, route similarity increased with time in the regular pentagonal array (Fig. 2B; GLMM_route similarity_: *Est*. = 0.110 ± 0.020, *P* < 0.001). When comparing simulations to experimental data, route similarity increased significantly more in model 1 (Fig. 2B; pos. reinf.; GLMM_route similarity_: *Est*. = 0.088 ± 0.020, *P* < 0.001) and in model 3 (Fig. 2B; pos. + neg. reinf.; GLMM_route similarity_: *Est*. = 0.086 ± 0.020, *P* < 0.001) than for real bees. However, route similarity in model 2 was significantly lower than for real bees (Fig. 2B; neg. reinf.; GLMM_route similarity_: *Est*. = −0.109 ± 0.020, *P* < 0.001). Similar trends were observed in the narrow pentagonal array (Fig. S5 and see Appendix S3).

Thus overall, positive reinforcement is necessary and sufficient to replicate the behavioural observations (although with a significant difference found for route similarity between the experimental data and the models 1 and 3), while negative reinforcement has no detectable effect.

### Simulations with two foragers

Having tested our models with one forager, we next explored conditions for the emergence of resource partitioning within pairs of foragers. Here experimental data are not available for comparison. We thus simulated foraging patterns and interactions of bees in different types of environments defined by flower patchiness. Each environment contained 10 flowers that were either distributed in one patch, two patches, or three patches (see examples in Fig. S2; for details, see Methods). Each bee had to visit five rewarding flowers to fill its crop to capacity.

### Exploitation and interference competition

We first analysed exploitation competition by quantifying the frequency of visits to non-rewarding flowers by each bee during each foraging bout. The frequency of visits to non-rewarding flowers decreased for simulated bees in model 2 (neg. reinf.; Fig. 3A; GLMM_1patch_: *Est*. = −3.32e-03 ± 1.90e-04; GLMM_2patches_: *Est*. = −2.10e-02 ± 2.00e-04; GLMM_3patches_: *Est*. = −2.06e-02 ± 2.00e-04) and model 3 (pos. + neg. reinf.; Fig. 3A; GLMM_1patch_: *Est*. = −8.94e-03 ± 2.20e-04; GLMM_2patches_: *Est*. = −1.88e-02 ± 3.00e-04; GLMM_3patches_: *Est*. = −1.05e-02 ± 2.00e-04), irrespective of the environment they were tested in. However, in model 1 (pos. reinf.), bees behaved differently in the different environments. In the one patch environment, bees decreased their visits to non-rewarding flowers (Fig. 3A; GLMM_1patch_: *Est*. = −4.26e-03 ± 2.10e-04), whereas in the two and three patch environments, bees tended to increase their visits to non-rewarding flowers (Fig. 3A; GLMM_2patches_: *Est*. = 6.27e-03 ± 1.80e-04; GLMM_3patches_: *Est*. = 6.65e-03 ± 1.90e-04).

**Figure 3.**
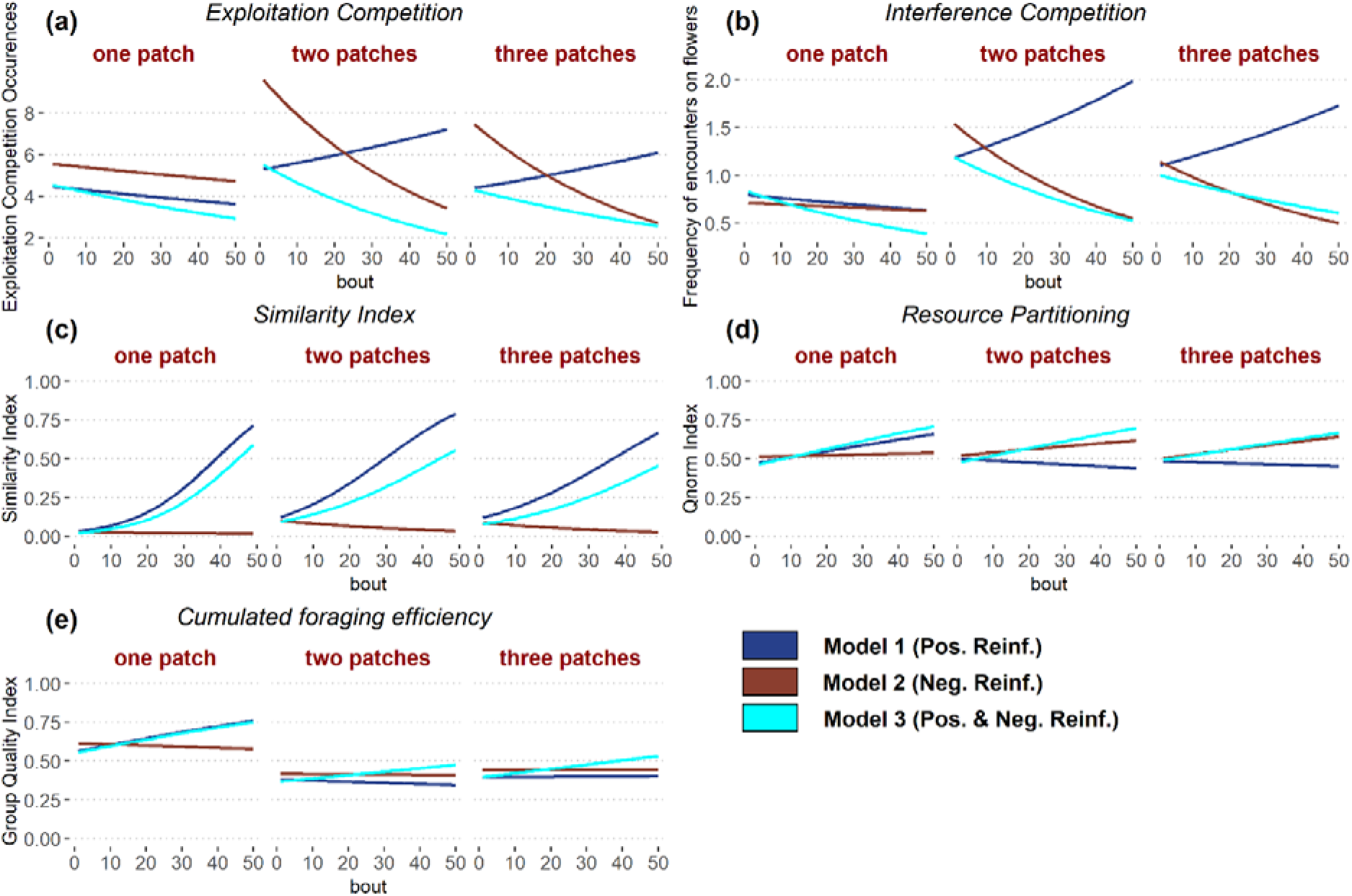
Results of simulations with two foragers in environments with 10 flowers. See details of models in Fig. 1. The x axis is the number of completed foraging bouts by the two foragers. The y axis represents respectively: (a) the estimated mean frequency of visits to empty flowers; (b) the estimated mean frequency of encounters on flowers; (c) the similarity index *SI_ab_* between two successive flower visitation sequences; (d) the index of resource partitioning *Q_norm_* (0: both bees visit the same set of flowers; 1: bees do not visit any flower in common); (e) the collective foraging efficiency index *QL_group_*. Average trends for each model are estimated across all types of environments (one patch, two patches and three patches; see Fig S2).

We analysed interference competition by quantifying the number of interactions on flowers at each foraging bout between the two bees. The frequency of encounters on flowers decreased with time for both model 2 (neg. reinf.; Fig. 3B; GLMM_1patch_: *Est*. = −2.49e-03 ± 7.40e-04; GLMM_2patches_: *Est*. = −2.10e-02 ± 6.00e-04; GLMM_3atches_: *Est*. = −1.68e-02 ± 7.00e-04) and model 3 (pos. + neg. reinf.; Fig. 3B; GLMM_1patch_: *Est*. = −1.53e-02 ± 8.00e-04; GLMM_2patches_: *Est*. = −1.66e-02 ± 7.00e-04; GLMM_3patches_: *Est*. = −1.01e-02 ± 6.00e-04), irrespective of the type of environment. Here again, bees of model 1 (pos. reinf.) behaved differently in the different environments. In the one patch environment, bees decreased their frequency of encounters on flowers (Fig. 3B; GLMM_1patch_: *Est*. = −4.57e-03 ± 7.20e-04), whereas in the two and three patches environments, bees increased their frequency of interactions (Fig. 3B; GLMM_2patches_: *Est*. = 1.05e-02 ± 4.00e-04; GLMM_3patches_: *Est*. = 9.16e-03 ± 5.20e-04).

Thus overall, negative reinforcement was necessary for reducing exploitation and interference competitions. By allowing bees to avoid empty flowers, negative reinforcement facilitated the discovery of new flowers and thus gradually relaxed competition. In the absence of negative reinforcement, both types of competition increased in environments with several flower patches.

### Route similarity

We analysed the tendency of bees to develop repeated routes by comparing the similarity between flower visitation sequences of consecutive foraging bouts for each individual (Fig. 3C).

Bees increased route similarity through time in all types of environments in model 1 (pos. reinf.; Fig. 3C; GLMM_1patch_: *Est*. = 1.34e-01 ± 2.00-e03; GLMM_2patches_: *Est*. = 9.56e-02 ± 1.30e-03; GLMM_3patches_: *Est*. = 7.76e-02 ± 1.20e-03) and model 3 (pos. + neg. reinf.; Fig. 3C; GLMM_1patch_: *Est*. = 1.33e-01 ± 2.00e-03; GLMM_2patches_: *Est*. = 6.95e-02 ± 1.20e-03; GLMM_3patches_: *Est*. = 6.14e-02 ± 1.3e-03). On the contrary, in model 2 (neg. reinf.), route similarity did not vary in the one patch environment (Fig. 3C; GLMM_1patch_: *Est*. = 7.46e-04 ± 6.65e-03) and decreased through time in the other environments (Fig. 3C; GLMM_2patches_: *Est*. = −1.91e-02 ± 2.90e-03; GLMM_3patches_: *Est*. = −3.20e-02 ± 3.10e-03).

The presence of negative reinforcement in models 2 and 3 reduced the final level of route similarity compared to trends found in model 1. In these conditions bees learned to avoid revisits to empty flowers and showed greater variation in their visitation sequences, as a result of searching for new flowers.

### Resource partitioning

We analysed the level of resource partitioning by quantifying the tendency of the two bees to use different flowers. This index varies between 0 (the two bees use the same set of flowers) and 1 (the two bees use completely different sets of flowers; see Methods).

In model 1 (pos. reinf.), bees showed an increase of resource partitioning with time in environments with one patch (Fig. 3D; GLMM_1patch_: *Est*. = 2.90e-02 ± 1.30e-03), and a decrease in environments with two or three patches (Fig. 3D; GLMM_2patches_: *Est*. = −1.02e-02 ± 1.30e-03; GLMM_3patches_: *Est*. = −8.26e-03 ± 1.26e-03). By contrast, in model 2 (neg. reinf.) and model 3 (pos. + neg. reinf.), bees showed an increase of resource partitioning with time in all types of environments (Fig. 3D; model 2: GLMM_1patch_: *Est*. = 1.22e-02 ± 1.30e-03; GLMM_2patches_: *Est*. = 1.28e-02 ± 1.20e-03; GLMM_3patches_: *Est*. = 1.82e-02 ± 1.20e-03; model 3: GLMM_1patch_: *Est*. = 3.55e-02 ± 1.30e-03; GLMM_2patches_: *Est*. = 3.17e-02 ± 1.30e-03; GLMM_3patches_: *Est*. = 2.19e-02 ± 1.30e-03). Model 3 displayed similar levels of partitioning in all the different environments where models 1 and 2 showed a greater variance. Model 1 had greater partitioning only in the one patch environment, while model 2 had greater partitioning in the two and three patch environments.

This suggests positive and negative reinforcements most likely contributed unevenly but complementarity in the model 3 with different spatial distributions of flowers. Positive reinforcement would be the main driver for partitioning in the one patch environment, while negative reinforcement would be the main driver in the two and three patches environments.

### Collective foraging efficiency

To quantify the collective foraging efficiency of bees, we analysed the capacity of the two foragers to reach the most efficient combination of route qualities (i.e. minimum distance travelled by a pair of bees needed to visit the 10 flowers; see Methods).

In model 1 (pos. reinf.), pairs of bees increased their collective foraging efficiency with time in environments of one and three patches (Fig. 3E; GLMM_1patch_: *Est*. = 4.20e-02 ± 1.50e-03; GLMM_3patches_: *Est*. = 3.04e-03 ± 1.25e-03). By contrast, bees decreased their level of foraging efficiency in the environment with two patches (Fig. 3E; GLMM_2patches_: *Est*. = −4.61e-03 ± 1.27e-03). In model 2 (neg. reinf.) pairs of bees decreased their collective foraging efficiency with time in all types of environments (Fig. 3E; GLMM_1patch_: *Est*. = −5.08e-03 ± 1.24e-03; GLMM_2patches_: *Est*. = −8.03e-03 ± 1.27e-03; GLMM_3patches_: *Est*. = −4.24e-03 ± 1.24e-03). In model 3 (pos. + neg. reinf.) bees increased their collective foraging efficiency with time in all types of environments (Fig. 3E; GLMM_1patch_: *Est*. = 4.12e-02 ± 1.50e-03; GLMM_2patches_: *Est*. = 8.77e-03 ± 1.25e-03; GLMM_3patches_: *Est*. = 1.83e-02 ± 1.30e-03).

The positive reinforcement seems to be the main driver for collective foraging efficiency in the one patch environment. However, none of the positive and negative reinforcements alone managed to increase the foraging efficiency in the two and three patch environments. Only their interaction, as seen in the model 3, brought an increase in the collective foraging efficiency. Collective efficiency is generally higher in the one patch environment than in the two and three patches environments because the difference between the best possible path (for which the collective foraging efficiency is equal to 1) and a typical suboptimal path of a simulated bee is lower due to the absence of long inter-patch movements.

## Discussion

Central place foraging animals exploiting patchily distributed resources that replenish over time are expected to develop foraging routes (traplines) minimizing travel distances and interactions with competitors (Possingham 1989; Ohashi and Thompson 2005; Lihoreau *et al*. 2016). Here we developed cognitively plausible agent-based models of vector navigation to explore the behavioural mechanisms leading to resource partitioning between traplining bees. In the models, bees learn to develop routes as a consequence of feedback loops that modify the probabilities of moving between flowers. Simulations show that, in environments where resources are evenly distributed, bees can reach high levels of resource partitioning based on positive reinforcement only, but cannot based on negative reinforcement only. However, in environments with patchily distributed resources, both positive and negative reinforcements become necessary.

When foraging in uniformly distributed plant resources (one patch), it is easiest to encounter all the resources available as none of them is isolated far from any other (with thus a low probability of being reached). Consequently, two bees are very likely, over time, to learn non-overlapping foraging routes and show resource partitioning. However, in environments with non-uniformly distributed resources (two or three patches), the added spatial complexity can interfere with this process. The initial likelihood of moving between distant patches is relatively low. Thus, the sole implementation of positive reinforcement often does not allow bees to explore all possible patches, so that the paths of competing bees overlap and interfere within a subset of the available patches. Adding a negative reinforcement for movement vectors leading to unrewarded flowers increases aversion for these empty flowers, the spatial segregation of foraging paths between competing bees and the collective exploitation of all available patches, even if the initial probabilities of moving to distant patches are low. This interplay between the influences of positive and negative experiences at flowers on the spatial and competitive decisions of bees is in accordance with the behavioural observations that bees tend to return to rewarding flowers and avoid unrewarding flowers, either because flowers were found empty or because the bees were displaced by a competitor during a physical encounter (Lihoreau *et al*. 2016; Pasquaretta *et al*. 2019).

The need for a negative reinforcement to enhance discrimination between different options or stimuli is well-known in learning theory and behavioural studies (Beshers and Fewell 2001; Garrison *et al*. 2018; Kazakova *et al*. 2020). At the individual level, negative experiences modulate learning. For both honey bees and bumblebees, adding negative reinforcement to a learning paradigm (e.g. quinine or salt in sucrose solution) enhances fine scale colour discrimination (Avarguès-Weber *et al*. 2010) and performances in cognitive tasks requiring learning of rules of non-elemental associations (Giurfa 2004). The insect brain contains multiple distinct neuromodulatory systems that independently modulate negative and positive reinforcement (Schwaerzel *et al*. 2003) and the ability of bees to learn negative consequences is well-established (Vergoz *et al*. 2007). At the collective level, negative feedbacks are also known to modulate social and competitive interactions. This is especially notable in collective decisions making by groups of animals and robots (Sumpter 2010), where negative feedbacks enable individuals to make fast and flexible decisions in response to changing environments (Robinson *et al*. 2005; Seeley *et al*. 2012). Even so, the utility of negative reinforcement to enhance efficient route learning and the consequences of this for the emergence of effective resource partitioning has not been commented on previously, it may be that this is a general phenomenon with applicability to other resource partitioning systems.

Our study implies that some very basic learning and interaction rules are sufficient for trapline formation and resource partitioning to emerge in bee populations, providing a solid basis for future experimental work. Nonetheless, several improvements of the model could already be considered for further theoretical investigations of bee spatial foraging movements and interactions, by implementing documented inter-individual variability in cognitive abilities (Chittka *et al*. 2003; Raine and Chittka 2012) and spatial strategies (Klein *et al*. 2017) of bees, the variability in the nutritional quality of resources (Wright *et al*. 2018; Hendriksma *et al*. 2019) and the specific needs of each colony (Kraus *et al*. 2019), or the well-known ability of bees to use chemical (Leadbeater and Chittka 2005) and visual (Dunlap *et al*. 2016) social information to decide whether to visit or avoid flowers. For instance, it is well-known that foragers of many bee species leave chemical cues as footprints on the flowers they have visited (bumblebees and honeybees: Stout and Goulson 2001; solitary bees: Yokoi and Fujisaki 2009). Bees learn to associate the presence or absence of a reward to these footprints and to revisit or avoid scented flowers (Leadbeater and Chittka 2011). Such a pheromone system is an advantageous signal for all participants in the interaction (Stout and Goulson 2001). This additional information could significantly enhance the positive or negative experiences of bees visiting flowers and thus increase resource partitioning to the benefit of all bees coexisting in the patch (Appendix S4), even of different species that learn to use these cues (Stout and Goulson 2001; Dawson and Chittka 2013). More exploration could also be done in the future in regards to the probability of winning a competitive interaction on flower. While we considered all individuals to have similar probabilities to access floral nectar when bees encounter on flowers, resource partitioning has been suggested to be favoured by asymmetries in foraging experiences (Ohashi et al. 2008; Lihoreau *et al*. 2016). Differences in experience or motivation would ultimately affect the outcome of competition, both passively (more consistent depletion of the flowers in a trapline) and actively (active displacement of other bees from one’s established trapline).

Our study fills a major gap of our understanding of pollinator behaviour and interactions by building on recent attempts to simulate trapline foraging by individual bees (Lihoreau *et al*. 2012b, Reynolds *et al*. 2013; LeMoël *et al*. 2019). As such, it constitutes a unique theoretical modelling framework to explore the spatial foraging movements and interactions of bees in ecologically relevant conditions within and between plant patches, thereby holding considerable premises to guide novel experiments. Further developments of the model could be used to test predictions with more than two bees (see examples Video S1 and Appendix S4), several colonies, or even different species of bees (e.g. honey bees) and thus complement current predictions about pollinator population dynamics (Becher *et al*. 2014; 2016; 2018). Ultimately, the robust predictions of the spatial movements and interactions of bees over large spatio-temporal scales, through experimental validations of the model, have the potential to show the influence of bee movements on plant reproduction patterns and pollination efficiency (Ohashi and Thomson 2009; Pasquaretta et al. 2017).

## METHODS

### Description of models

We built three agent-based models in which bees learn to develop routes in an array of flowers (see summary diagram in Fig. 1). The environment contains flowers each delivering the same quality and volume of nectar. At each foraging bout (flower visitation sequence, beginning and ending at the colony nest entrance as the bee starts and finishes a foraging trip, respectively), each bee attempts to collect nectar from five different flowers in order to fill its nectar crop (stomach) to capacity. Flowers are refilled between foraging bouts. In simulations with two bees, the two individuals start their foraging bout synchronously and the flowers are refilled with nectar after the last bee has returned to the nest. For each bee, flower choice is described using movement vectors (orientated jump between two flowers or between the nest and a flower). The initial probability of using each possible movement vector is based on the length of the movement vectors, so that short vectors have a higher probability than longer ones. This probability is then modulated through learning when the bee used a vector for the first time during a bout.

We implemented two learning rules: (i) a positive reinforcement, i.e. if the flower at the end of a movement vector contains nectar and the bee feeds on it, it is set as a rewarding experience and the probability to reuse the vector later is increased; (ii) a negative reinforcement, i.e. if the flower is empty or if the bee is pushed away by competitors, it is set as a non-rewarding experience and the probability to reuse the vector later is decreased. The three models implemented either one of these two rules (model 1: positive reinforcement only; model 2: negative reinforcement only) or both rules (model 3).

A flower is empty if it had previously been visited in the same foraging bout by the same or another bee (exploitation competition). If multiple bees visit a flower at the same time (interference competition), only one bee (randomly selected) is allowed to stay and take the reward if there is one. The other bees react as if the flower is empty. After each flower visit, all bees update their probabilities to reuse the vector accordingly.

Route learning thus depends on the experience of the bee and its interactions with other foragers. For simplicity, we restricted our analysis to two bees. Working with pairs of bees facilitates future experimental tests of the models’ predictions by reducing the number of bees to manipulate and control in experiments (Ohashi et al. 2008; Lihoreau et al. 2016). Note however that the same models can be used to simulate interactions among more bees (see examples with five bees in Video S1, Appendix S4).

A detailed description of the model is provided in Appendix S1, in the form of an Overview, Design concepts and Details (ODD) protocol (Grimm et al. 2006; Grimm et al. 2020). The complete R code is available at *https://gitlab.com/jgautrais/resourcepartitioninginbees/-/releases*.

### Environments

#### Simulations with one forager

Our first goal was to test the ability of our models to replicate observations of real bees. To do so, we ran simulations in environments replicating published experimental studies in which individual bumblebees (*Bombus terrestris*) were observed developing traplines between five equally rewarding artificial flowers in a large open field (Lihoreau *et al*. 2012b; Woodgate *et al*. 2017). To our knowledge, these studies provide the most detailed available datasets on trapline formation by bees. Lihoreau *et al*. (2012b) used a regular pentagonal array of flowers (Fig. S1A) in which they tracked seven bumblebees. We judged this sample size enough to run quantitative comparisons with model simulations (raw data are available in Table S1 of Lihoreau *et al*. 2012b). Woodgate *et al*. (2017) used a narrow pentagonal array of flowers (Fig. S1B). Here, however, the small sample size of the original dataset (three bumblebees, data shared by J. Woodgate) only enabled a qualitative comparison with the model simulations (Appendix S3).

#### Simulations with two foragers

We then explored conditions leading to resource partitioning by running model simulations with two foragers. Here we simulated environments containing 10 flowers, in which each bee had to visit five rewarding flowers to fill its crop to capacity. To test whether model predictions were robust to variations in spatial distributions of resources, we simulated three types of environments characterized by different levels of resource patchiness: (*i*) a patch of 10 flowers, (*ii*) two patches of five flowers each, and (*iii*) three patches of five, three and two flowers respectively (see examples in Fig. S2). We generated flower patches into a spatial configuration comparable to the one used in both experimental setups (Lihoreau *et al*. 2012b; Woodgate *et al*. 2017). In a 500m*500m plane, a nest was set as the centre (coordinates 0;0). Then, patch centres were placed with a minimum distance of 160m between each, and at least 20m from the nest. Within a patch, flowers were randomly distributed according to two constraints: (i) flowers were at least 20m apart from each other and from the nest, (ii) the maximum distance of each flower from the centre of their patch was 40m. This ensured that each patch had a maximum diameter of 80m and inter-flower distances were smaller between all flowers of the same patch than between all flowers of different patches (See ODD Protocol for more details, Appendix 1, Ch.7 “Submodels”).

### Movements

At each step, a bee chooses to visit a target location (flower or nest) based on a matrix of movement probabilities. This matrix is initially defined using the inverse of the square distance between the current position of the bee and all possible target locations (Lihoreau *et al*. 2012b; Reynolds *et al*. 2013). The probability of moving from location *i* to the location *j* among *n* possible targets, is initially set to:

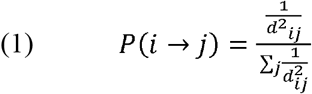

Where *d_ij_* is the distance between locations *i* and *j*. Before choosing its destination, the bee lists all possible target locations. For simplicity, the bee excludes its current location, thus preventing looping flights to and from the same target (flower or nest), which are rare in experienced bumblebee foragers (Saleh and Chittka 2007) and provide little information about bee routing behaviour. The bee also excludes the location it had just come from to simulate the tendency of bumblebees to avoid recently visited (and thus depleted) flowers (Saleh and Chittka 2007). The foraging bout ends if: (*i*) the bee fills its crop to capacity, (*ii*) the bee chooses the nest as a target destination, or (*iii*) the bee reaches a maximum travelled distance of 3000 m. The latest was added to avoid endless foraging trips in the model. The maximum distance was chosen based on the observation that bumblebees typically forage within a distance of less than 1km from their nest (Osborne *et al*. 1999; Wolf and Moritz 2008; Woodgate *et al*. 2016).

### Learning

Learning modulates the probability of using movement vectors as soon as the bee experiences the chosen target and only once within a foraging bout (the first time the movement vector is used this bout; Fig. 1). This approach has the advantage of implementing vector navigation (Stone *et al*. 2017; Le Moël *et al*. 2019) and thus avoids assumptions about computation and comparison of complete routes (Lihoreau *et al*. 2012b; Reynolds *et al*. 2013). Positive reinforcement was implemented in models 1 and 3. It occurred when a bee used a vector leading to a rewarding flower. The probability of using this vector was then multiplied by 1.5 (other vectors probabilities were scaled accordingly to ensure that all sum up to 1) as in Reynolds *et al*. (2013). This positive reinforcement is based on the well-known tendency of bumblebees to return to nectarrewarding places through appetitive learning (Goulson 2010). Negative reinforcement was implemented in models 2 and 3. It occurred when a bee used a vector leading to a non-rewarding flower. The bee reduced the probability of using that vector by multiplying it by 0.75 (here also rescaling the probabilities after application of the reinforcement). This negative reinforcement rule was based on the tendency of bumblebees to reduce their frequency of revisits to unrewarded flowers with experience (Pasquaretta *et al*. 2019). We applied a lower value to negative reinforcement because bees are much more effective at learning positive stimuli (visits to rewarding flowers) than negative stimuli (visits to non-rewarding flowers) (review in Menzel 1990). Sensitivity analyses of these two parameters show that increasing positive and/or negative reinforcement increases the speed and level of resource partitioning (see Appendix S4).

### Competitive interactions

We implemented competitive interactions between foragers in the form of exploitation and interference (Fig. 1). Exploitation competition occurred when a bee landed on a flower whose nectar reward had already been collected by another bee. If the flower was empty, the probability to reuse the vector was either left unchanged (model 1) or decreased (negative reinforcement, models 2 and 3). Interference competition occurred when two bees encountered on a flower. Only one bee could stay on the flower and access the potential nectar reward with a random probability (p=0.5). After the interaction, the winner bee took the reward if there was one. The loser bee reacted as it would for an empty flower.

### Data Analyses

All analyses were performed in R version 3.3 (R Core Team 2018).

#### Simulations with one forager

For each model, we compared the results of the simulations to the reference observational data, either quantitatively (for Lihoreau *et al*. 2012b) or qualitatively (for Woodgate *et al*. 2017; see Appendix S3). We stopped the simulations after the bees completed a number of foraging bouts matching the maximum observed during the experimental conditions of the published data (37 foraging bouts in Lihoreau *et al*. 2012b; 61 foraging bouts in Woodgate *et al*. 2017). We ran 500 simulations for each model and we estimated how models fitted the experimental data using two main measures:

i. the quality of each route, *QL*, calculated as:

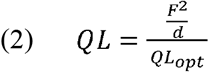 Where *F* is the number of rewarding flowers visited during a foraging bout and *d* is the net length of all vectors travelled during the foraging bout. *QL* is standardized in [0; 1] by the quality of the optimal route in each array *QL_opt_* (shortest possible route to visit all 5 flowers).
ii. a similarity index *SI_ab_* between flower visitation sequences experienced during two consecutive foraging bouts *a* and *b* as follows:

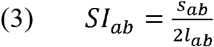

Where *s_ab_* represents the number of flowers in vectors found in both sequences, and *l_ab_* the length of the longest flower visitation sequence between *i* and *j* multiplied by 2 to make sure that *SI_ab_* = 1 occurs only when two consecutive sequences sharing the same vectors also have the same length (more details and examples in Appendix S5).

We applied generalized linear mixed effect models (GLMM) with binomial error, using the *glmer* function in ‘lme4’ package (Bates *et al*. 2015), to assess whether the estimated trends across foraging bouts for *QL* and *SI_ab_* obtained from model simulations with one forager differed from trends obtained from experimental data. In each model, we used a random structure to account for the identity of bees.

#### Simulations with two foragers

We generated 10 arrays of flowers for each of the three types of environments (one patch, two patches and three patches) and ran 100 simulations for each of the three models (9000 simulations in total). We compared the simulation outcomes of the models using four measures:

i. the frequency at which each bee experienced exploitation competition (i.e. flower visits when the reward has already been collected) and interference competition (i.e. flower visits when two bees encounter on the flower).
ii. the similarity index *SI_ab_* between successive foraging bouts by the same bee.
iii. the degree of resource partitioning among bees, based on network modularity *Q* (Pasquaretta and Jeanson 2018; Pasquaretta *et al*. 2019). *Q* is calculated using the *computeModules* function implemented in the R package ‘bipartite’ (Dormann *et al*. 2008) using the *DIRTLPAwb+* algorithm developed by Beckett (2016). Although *Q* ranges between 0 (the two bees visit the same flowers) and 1 (the two bees do not visit any flower in common), the comparison of modularity between networks requires normalisation because the magnitude of modularity depends on network configuration (e.g., total number of flower visits) (Dormann and Strauss 2014; Beckett 2016). For each network, we calculated:

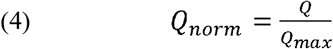

where *Q_max_* is the modularity in a rearranged network that maximizes the number of modules (Pasquaretta and Jeanson 2018).
iv. an index of collective foraging efficiency, *QL_group_*, computed for each foraging bout *b*, to estimate the collective efficiency of all foraging bees, as:

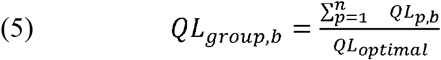

where *QL_p,b_* is the route quality of the individual *p* during bout *b, n* the number of bees, and *QL_optimal_* is the maximum value of all the possible sums of individual route qualities. *QL_optimal_* was calculated in each environment by computing all possible combinations of two routes visiting five flowers each and extracting the combination with the highest quality.

To assess whether the trends across foraging bouts obtained from simulations with two bees differed between models (Fig. 1) and types of environments (Fig. S2), we applied GLMMs for each of the following response variables: (*i*) frequency of competition types (Poisson error distribution), (*ii*) *SI_ab_* (Binomial error distribution), (*iii*) *Q_norm_* (Binomial error distribution) and (*iv*) *QL_group_* (Binomial error distribution). In each model, we used a random structure to account for bee identity nested in flower arrays (i.e. 100 simulations of each spatial array for each model). To statistically compare the trends across foraging bouts, we estimated the marginal trends of the models, as well as the 95% confidence intervals of the mean using the *emtrends* function in ‘emmeans’ package (Lenth 2018). When the 95% confidence intervals of the estimated trends included zero, the null hypothesis was not rejected. Statistical models were run using the *glmer* function in ‘lme4’ package (Bates *et al*. 2015).

## Statement of authorship

**Thibault Dubois**: Conceptualization, Software, Formal Analysis, Writing – Original Draft, Writing – Review & Editing. **Cristian Pasquaretta**: Conceptualization, Formal Analysis, Writing – Original Draft, Writing – Review & Editing. **Andrew B. Barron**: Writing – Review & Edition. **Jacques Gautrais**: Software, Writing – Review & Editing. **Mathieu Lihoreau**: Conceptualization, Writing – Original Draft, Writing – Review & Editing, Supervision.

## Data Availability Statement

Simulation code is archived in GitLab: *https://gitlab.com/jgautrais/resourcepartitioninginbees/-/releases*.

## Acknowledgements

We thank Joe Woodgate for sharing raw flower visitation data of Woodgate *et al*. (2017). We are also grateful to Matthias Becher and Mickaël Henry for useful comments on a previous version of this manuscript.

## Supplementary materials

## Appendix S1. ODD Protocol

Below we provide a description of our models following the ODD (Overview, Design concepts, Details) protocol (Grimm et al. 2006, 2020).

### 1 Purpose and patterns

The purpose of the models is to offer an explanation to how multiple bees learn to optimize their foraging efficiency. Bees are expected to do so by developing efficient routes (traplines) minimizing spatial overlaps with other foragers (resource partitioning) (Lihoreau et al., 2016). We suggest such process can be achieved through combinations of positive and negative reinforcements during flower visits.

### 2 Entities, state variables and scales

The models depict three kinds of entities: bees, flowers and the colony nest. Bees go out foraging for nectar rewards on flowers and return to the nest. Numbers of bees and flowers can be adjusted in the model. In our simulations we set a ratio of five flowers per bee as to fit the experimental data we replicated (Lihoreau et al., 2012; Woodgate et al., 2016). A single colony nest is represented, from which all bees forage and come back to. The bees are defined by the following set of state variables:

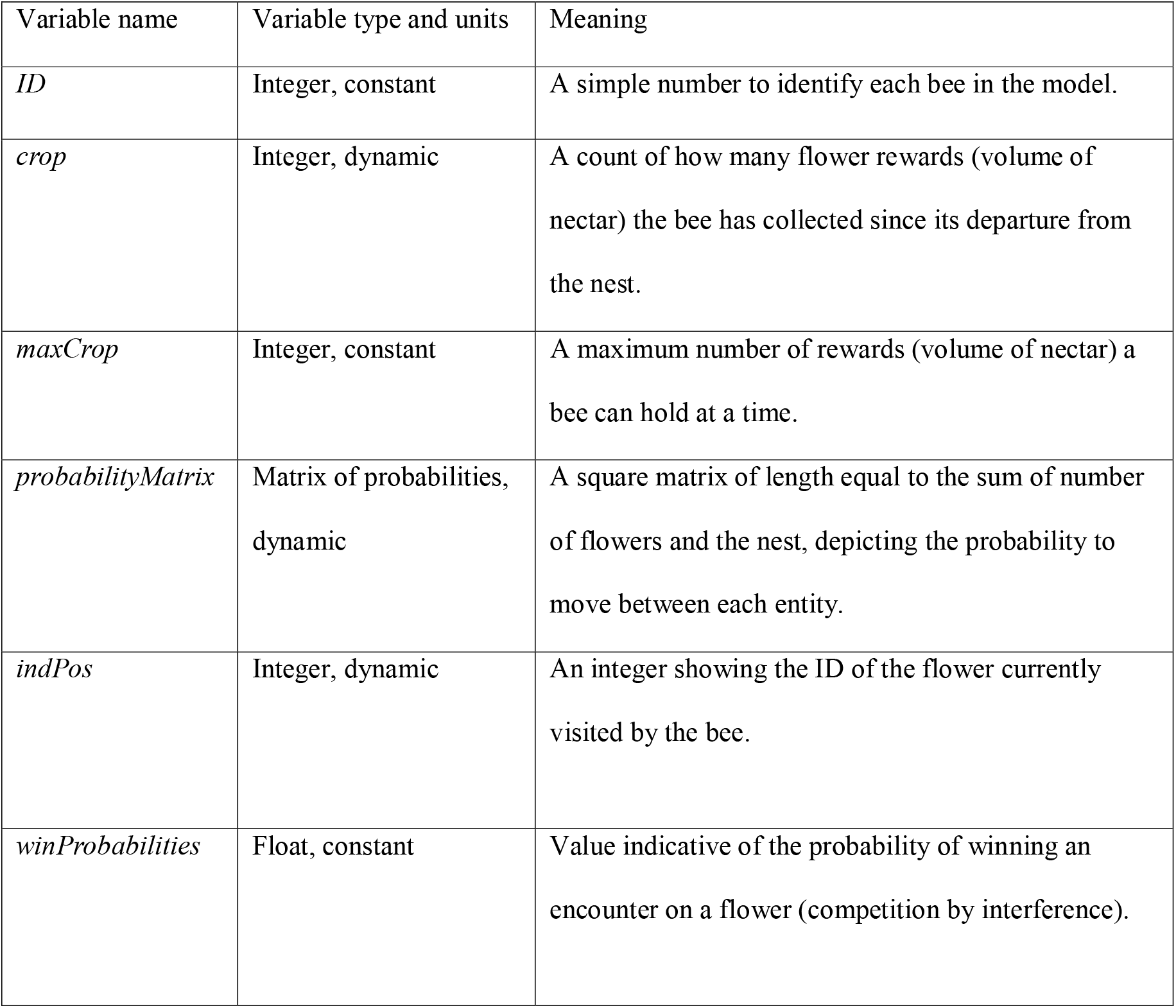

The flowers are static entities placed in the environment. They are defined by:

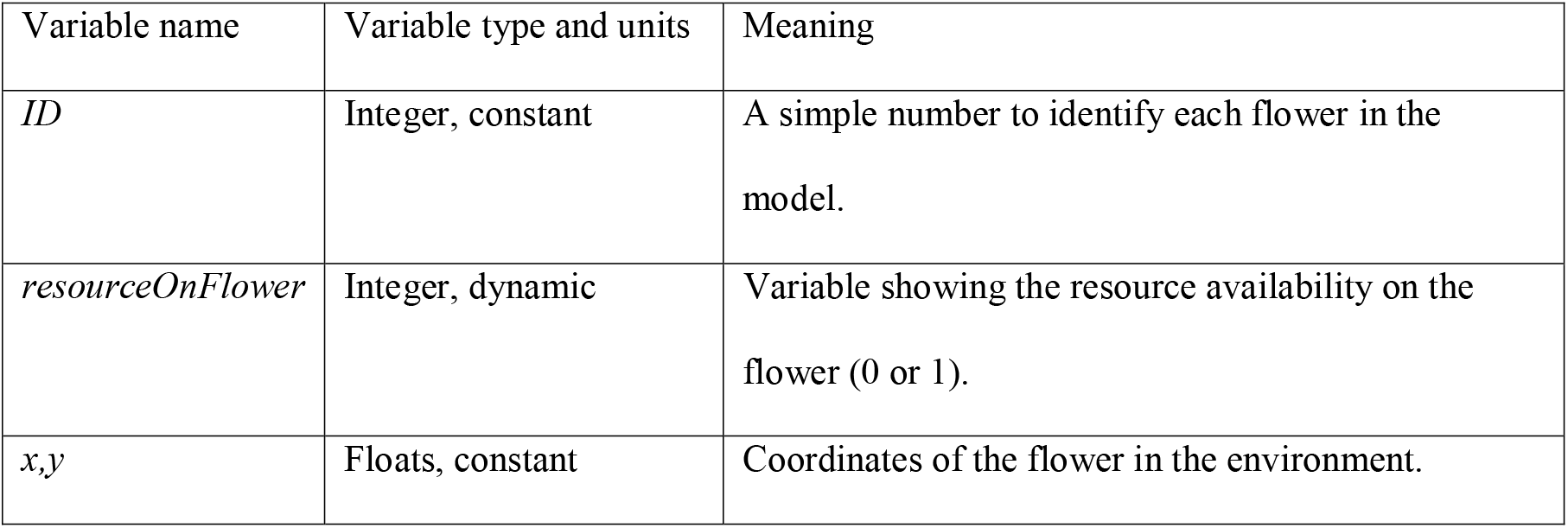

The nest is a unique entity which is represented by the following variables:

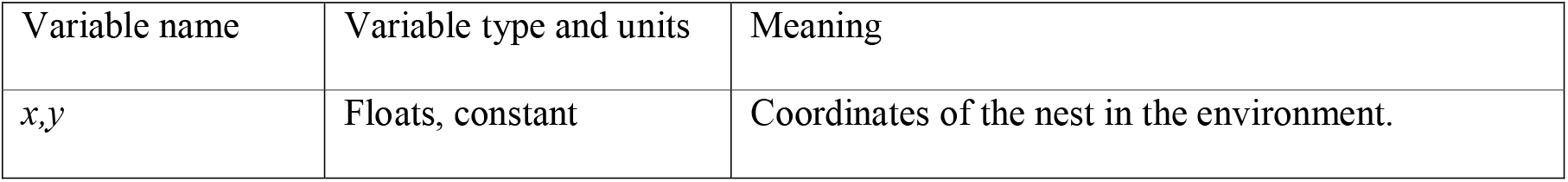

Both the spatial and temporal scales are represented. Space is represented by the relative distance between the different flowers. Internal parameters define min/max ranges in which each entity can be placed, relative to other entities (see *CreateEnvironment* submodel). Time is represented abstractly. At each step, the bees visit a flower, and possibly feed on it. We assume all travel and flower manipulation times are identical.

### 3 Process overview and scheduling

The model covers the execution of a series of foraging bouts (foraging trip starting and ending at the colony nest) by all the bees. During a foraging bout, the model keeps running as long as one bee is foraging. The number of foraging bouts performed by the bees is set as a parameter of the simulation. Each foraging bee chooses its next destination, using the *ChooseDestination* submodel, and updates its position to this new destination. This action is executed in the order of the bees’ *ID* value. If a bee chooses the nest as next destination, its bout is over. Before feeding, the bees resolve any competition occurrences (if more than one bee is on a single flower), in the order of the flower’s *ID* on which the competition occurs. A weighed sampling of the *winProbabilities* decides which individual wins the interaction (default values are uniform for all bees). Bees that either were alone on a flower or won an interaction feed on the flowers, using the *Feeding* submodel. All foraging bees update their movement probability matrix according to their recent experience (details in the *Learning* submodel). Finally, each foraging bee checks if it has reached one of the conditions to return to the nest (crop full or maximum distance of travel), and if so, finishes its foraging bout.

### 4 Design concepts

#### a Basic principles

Animals are expected to distribute themselves among different resources to optimize the energetic intake, following what is commonly called the Ideal Free Distribution theory (Fretwell & Lucas, 1969). The environment in which bees forage present multiple constraints. Flowers provide a renewable resource in very low amounts and have a fast turnover changing the distribution of resources. Competition with other bees is strong but also very dynamic as new foragers arrive daily while older ones die.

Bees tend to optimize their foraging activity (Ohashi et al., 2007; Lihoreau et al., 2012b; Lihoreau et al., 2016) by traplining (developing stable routes between feeding sites) and partitioning resources (exploiting feeding sites in different areas of the environment to minimize competition). The aim of the model is to explore the conditions under which traplining emerges from the behaviour of independent bees interacting with the resources and other agents according to the principles of the Ideal Free Distribution. The models reuse parts of a previously published model (Lihoreau et al. 2012b; Reynolds et al., 2013) which has been to our knowledge the only model trying to represent the ontogeny of the traplining behaviour to this date.

#### b Emergence

We focused on how both optimization strategies (traplining and partitioning) can emerge from simple rules of positive and negative reinforcements derived from the foraging experience of individual bees. We explored how these outcomes change when we alter the spatial distribution of the flowers, or remove either type of reinforcement. When bees partition resources, the partitioning index Qnorm (Pasquaretta & Jeanson, 2018) reaches a maximum value of 1. When bees follow traplines, the cumulated foraging efficiency and route similarity index of the bees approaches a maximum value of 1.

Resource partitioning and traplining are emergent and vary depending on the parameters of the simulation. While traplining could be achieved with a positive reinforcement only, resource partitioning required both positive and negative reinforcements to emerge.

#### c Adaptation

Bees follow a heuristic in the form of a matrix of movement probability to choose their next destination. This matrix changes with experience to favour the flowers where the bees had a positive experience (collection of nectar). Bees thus learn to fill their nectar crop to capacity. More information on the probability matrix is given in the *ChooseDestination* submodel.

#### d Objectives

The bees have a unique goal: finding a set amount of nectar to fill their crop (whose capacity is set by the *maxCrop* parameter). In all the conditions explored, this crop capacity was set to 5 resources (units of nectar volume): an arbitrary value used to fit the experimental data of Lihoreau et al. (2012b). The decision-making process is altered by this crop filling (described by the *crop* variable) if it reaches the same value as *maxCrop*. If so, the foraging bout is over, and the bee returns to the nest. This behaviour has been observed in experimental conditions (Lihoreau et al., 2012b; Woodgate et al., 2016).

#### e Learning

The learning process occurs as changes made to the probability matrix during the simulation through two components: a positive and a negative reinforcement.

Positive reinforcement is triggered when a bee finds a resource on a flower. When it happens, the bee increases the probability of reusing the vector it just used. Values of the positive reinforcement factor are typically set to be superior or equal to 1. Negative reinforcement is similar but reduces the probability of reuse of the vector. Negative reinforcement occurs when there is either no resource on the flower or if the bee has been evicted from the flower by a competitor. Values of the negative reinforcement factor are typically set to be less than or equal to 1.

#### f Prediction

In the models the movement probability matrix acts as a prediction tool for the bees. After some experience in the environment, these probabilities become proxies for the probability of finding nectar in flowers, as the positive and negative experiences shape the matrix. The choice of visiting a flower is done prior to the knowledge of presence of a reward on the flower, and only relies on the previous experience on this flower. More information on the movement probability matrix is given in the *ChooseDestination* submodel.

#### g Sensing

When the model is initialized, the movement probability matrix is based on the distances the bees have to travel between each flower. However, the model does not depict a probability of “not finding any flower”. In experimental situations, it is not rare that the bees would not find all the flowers during their first bout (Lihoreau et al., 2012; Woodgate et al., 2016). However, in our models it is assumed that the bee always finds a flower, even if it has never visited it. Moreover, it is assumed that the bee knows how to link all the flowers together (knowledge of all the existing vectors) even if these links have never been performed. The use of distance-weighed probabilities is a good approximation of the probabilities obtained by a random walk. As bees always keep tracks of their successive movements after leaving the nest, they can sum these movements to know the direction and approximate distance of the nest.

#### h Interactions

Two types of interactions were included: exploitation competition and interference competition. Exploitation competition occurs when a bee visits a flower that has already been depleted by a competitor or by itself during a prior visit. Interference competition occurs when two or more bees are simultaneously present on the same flower. When this direct interaction happens, only one bee can access the nectar reward. The winning bee is selected using uniform probabilities.

#### i Stochasticity

Stochasticity is included in three parts of the model: to place all the flowers in the environment, to choose a bee’s next destination, and to choose a winner from interference competition. When generating a new environment, the submodel *CreateEnvironment* sets rules for placing the flowers. The algorithm will repeatedly try to place a flower at a random position until it fits all the conditions.

The choice of the bee’s next destination (described in the *ChooseDestination* submodel) is done by consulting the movement probability matrix of the agent. As this choice relies on probabilities, we used a stochastic process to choose the next destination, according to the weights of the different destinations’ probabilities. The reason behind this choice is explained in the *Learning* and *Sensing* part of this ODD. Finally, the winner of an interference interaction is decided by choosing with uniform probabilities which bee wins.

#### j Collectives

The model includes no collectives.

#### k Observation

There are two observations collected from the bees: visitation sequences and occurrences of competition.

Every time a bee visits a flower during a foraging bout, the flower’s *ID* is reported into a vector containing all the visits in order during the bout. Every time a bee finds an empty flower or finds a competitor on the same flower, these occurrences are reported throughout the foraging bouts. This information gives the number of competitive interactions throughout the bouts. We also save the spatial positions of the flowers to compute the distances between them and find the theoretically optimal routes for the agents. This allows to compute the route qualities of the agents’ successive bouts.

### 5 Initialization

The first part of the initialization of the models is to create the environment. This part, described in details in the *CreateEnvironment* submodel, places all the flowers and the nest. The environment can be initialized in two ways: either by calling for an existing one (all the environments are stocked in a folder called *Arrays*, inside the main folder where the R script is), or by using the models to generate one.

All the parameters of the bees are then initialized. The *maxDistance* a bee can travel during a foraging bout, the *maxCrop* capacity of a bee’s nectar crop, the values of the *learningFactor* and *abandonFactor* which relate to the positive and negative reinforcements, respectively, and the *winProbabilities* in case of a direct interference competition. These values remain constant throughout the simulation. The variables of the bees that will be dynamic during the simulation are initialized: *crop*, the number of resources gathered during the ongoing foraging bout; *distanceTravelled* the distance travelled by the bee during this bout; and finally *boutFinished*, a Boolean indicating if the bee has finished its bout or not. The movement probability matrix is then initialized for each bee. Finally, we initialize the output objects to store the visitation sequences and resources gathered by each bee during each bout, and store all the information we inputted as parameters to be able to identify what were the parameters used for this simulation later.

While some simulations were made to be case-specific (emulating the experiment from Lihoreau et al. (2012b)) all the others were generic. For this specific case, we created our own files in a subfolder of the *Arrays* folder. This subfolder named *lihoreau2012* contained two files: *arrayGeometry.csv* and *arrayInfos.csv*. Their contents were as follow:

#### arrayGeometry

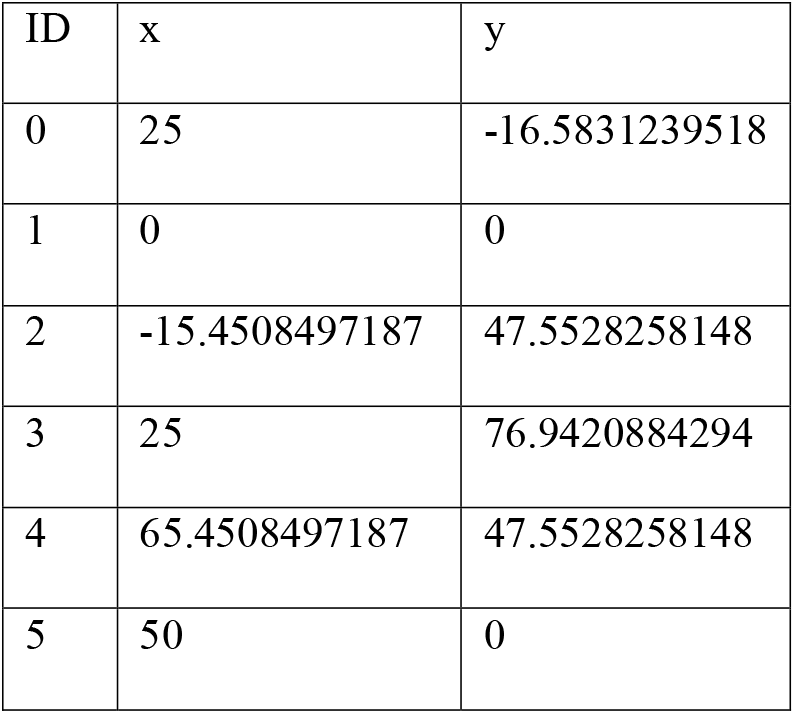

#### arrayInfos

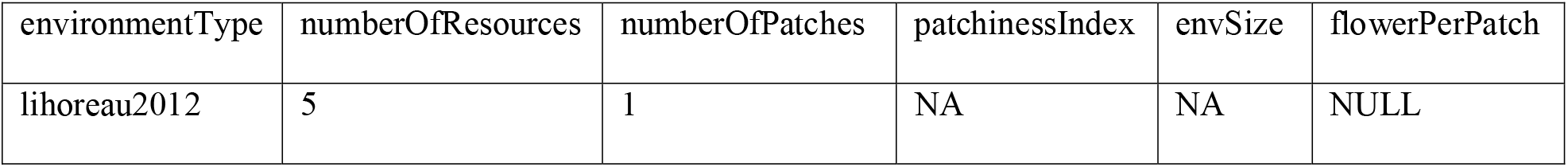

They are CSV files with a comma separator and a dot decimal. To fit with this case, the parameter *numberOfBees* had to be set to 1, as this was an experiment with a single bee. Also, the *environmentType* parameter in the script also has to be set to “*lihoreau2012*” in this case (same name as the subfolder and the *environmentType* value in the CSV file).

### 6 Input data

The model contains an R script dedicated to the input parameters. While all parameters have comments to help the user, a description of the important parameters is given here.

#### environmentType

Can either contain “generate” or the name of an existing and valid environment found in the *Arrays* folder (the *Arrays* folder will only be created after a first simulation, unless it is created manually beforehand). If using an existing environment, the parameters in part 3.1. will be ignored.

#### numberOfResources

Total number of flowers in the environment.

#### numberOfPatches

In how many patches these flowers should be distributed.

#### patchinessindex

Deprecated index, should be kept at 1. If multiple patches are created, a value of 1 will ensure the patches are set very distinctly from each other. Values closer to 0 will create a less defined limit between patches. Inputs between 0 and 1 are accepted.

#### envSize

The size of the environment. It defines a square environment in which the flowers will be placed. However, if it is impossible to place all the flowers, this value will be incrementally increased to give enough space.

#### flowerPerPatch

Should contain as many values as the number of patches. If only one patch, it must be set to NULL. Otherwise, it must take a succession of values following the syntax *c(a, b, c,…), a, b* and *c* being integer values whose sum equals to *numberOfResources*.

#### numberOfArrays

The number of different environments the model should generate using these parameters.

#### reuseGeneratedArrays

Can take TRUE or FALSE. If TRUE, the model will look into the *Arrays* folder for environments fitting all the parameters. If it does find similar ones, it will use them instead of generating new ones.

#### numberOfBees

Number of bee agents in the model.

#### numberOfSimulations

How many simulations the model will do for each set of parameters.

#### numberOfBouts

How many foraging bouts each bee will do during a simulation.

#### distFactor

Weight given to distance when generating the movement probability matrix. The probability for a vector of distance *d* is computed as probability = 1/*d^distFactor*. Changing this number will change the initial probability matrix.

#### param.useRouteCompare

deprecated, used to switch between our *Learning* submodel and the route comparison model of Lihoreau et al. (2012b) and Reynolds et al. (2013). Should be left at FALSE.

#### param.learningFactor

the value used for the positive reinforcement process. Values should be greater than or equal to 1. Requires at least one value. If multiple values are inputted, the model will run the simulations for each value.

#### param.abandonFactor

The value used for the negative reinforcement process. Values should be between 0 and 1.

#### maximumBoutDistance

Maximum distance a bee can travel during a foraging bout.

In the “Advanced parameters” category different rules can be enforced on the bees. In the following we detail the ones used in our simulations:

#### allowNestReturn

Allows the bee to select the nest as its next destination in the *ChooseDestination* submodel, based on the distance-weighed probabilities. If the bee does so, the foraging bout is finished.

#### forbidReverse Vector

This rule forbids the bees to use the reverse movement vector from the one they just used. If the bee has just moved from flower 2 to flower 3, for its next movement this bee will not be given the choice to go from flower 3 to flower 2. This interdiction only applies for the last vector executed.

#### onlineReinforcement

This rule forces the trigger of the *Learning* submodel after each encounter of a flower, instead of only triggering it when the bee had finished its bout. Movement probabilities are thus altered directly after the execution of a vector.

### 7 Submodels

Our models can be decomposed in multiple different submodels, which are described here.

#### CreateEnvironment

The creation of an environment happens first in the initialization of the model. The code relating to this process is found in the Functions script, in a function of the same name.

If the user chooses an *environmentType* different than “generate” the model will import the user’s selected environment. The creation of the environment using the “generate” option calls an algorithm we designed to create flower patches. It follows arbitrary rules without any ground in experimental data or theoretical background. In this function, all the parameters inputted in the 3.1.1 part of the Parameters script are being used. Refer to their description in the Input category for their meaning. The basic distance unit between entities is set by an internal parameter, *perceptionRange*, whose default value is 10.

The nest is set first at the centre of the environment (coordinates (0,0)). The different patch centres are placed between 2\**perceptionRange* and *envSize* from the nest. Every time a patch is placed, the algorithm checks if this patch centre is at least 16\**perceptionRange* away from any other patch centre. The algorithm reiterates this process until the condition is verified. The patch centres act as the first flower of each patch.

Flowers are then placed around the patch centres, respecting the distribution specified in *flowerPerPatch*. All flowers are tentatively placed between 2\**perceptionRange* and 4\**perceptionRange* of the patch centre, and must verify the condition that each flower has to be at least *2*perceptionRange* from any other flower. The algorithm reiterates this process until the condition is verified. If the algorithm fails to place a flower 200 times in a row, the range at which the flowers can be placed around the patch centre becomes between 2\**perceptionRange* and 4\**perceptionRange* + (*envSize*/20).

Once all the flowers are placed in all the patches, a table containing the flowers’ *ID*, coordinates *x* and *y*, and the patch they belong to (numbered from 1 to *numberOfPatches)* is created. The nest is also represented in this table, and takes the *ID* 0, and is part of its own patch.

#### ChooseDestination

In order to choose among the possible destinations, the bees refer to a movement probability matrix they are given at the beginning of the simulation. This matrix has *n* rows and columns, *n* being the number of entities (flowers and nest). It is created by extracting in a similar matrix the distance between each entity, and from then applying the following formula for each cell: Probability = 1 / d^*distFactor*

Where *d* is the distance indicated in the cells, and *distFactor* is an integer parameter, whose default value is 2. See the Input part for more information about *distFactor*. The probability to go from a flower to itself (immediate revisit) was set to 0. Visiting the same flower twice in a row happens when bees come back to the last departed flower if their search for another flower was unsuccessful, or if they do short orientation flights on the flower. However, these revisits have little importance for the establishment of a stable route (Lihoreau et al., 2010), and were thus ignored. All rows of the probability matrix are normalised so that their sum is equal to 1. To choose a bee’s next destination, it looks at the matrix’s row matching the flower ID of its current position. As the use of a “reverse vector” is forbidden (see the *forbidReverseVector* parameter described in the Input data part), the bee’s previous position is removed from the possible destinations. This prevents an artificial situation the model could create when two flowers are very close to each other, and the probability to move between them is much higher than any other probability. Without this rule, bees would often get stuck navigating back and forth between both flowers. If the *allowNestReturn* is used, the nest is kept in the possible destinations. Otherwise, it is removed. A weighed sample is made between all the remaining potential destinations to choose the one that the bee will use.

#### Feeding

This submodel takes care of all matters that happen when a bee lands on a flower, i.e. all competition occurrences and the collection of resources. If two or more bees choose the same destination on the same step, one of them is chosen randomly to access to the resource. The losing bees will still depart from this flower for the next step, but will not feed.

#### Learning

As bees finish to go through the *Feeding* module, they have five possible outcomes: (i) the bee has landed alone on a flower, and found resources; (ii) the bee has landed alone on a flower, and did not find any resource; (iii) the bee has landed with competitors on a flower, and has lost the competition; (iv) the bee has landed with competitors on a flower, has won the competition, but found no resource; (v) the bee has landed with competitors on a flower, has won the competition, and found resources. These outcomes can be placed into two categories: the positive (i and v) or negative (ii, iii and iv) outcomes.

Each bee has a square matrix named *indFlowerOutcome*, with *n* rows and columns, *n* being the combined number of flowers and nest. Every bout it is initialized, filled with 0s in all cells. After each outcome of a vector, the value of the cell corresponding to this vector is changed to 1 or −1 for a positive or negative experience, respectively. The probability linked to this same vector is then altered in the probability matrix, by multiplying the previous value by the appropriate reinforcement factor (*learningFactor* or *abandonFactor* for the positive and negative reinforcements, respectively). All probabilities in the altered row are then normalised to bring its sum back to 1.

## Appendix S2 Sensitivity analysis of positive and negative reinforcements

We ran a sensitivity analysis for the two main parameters: the positive and negative reinforcements. As we had no *a priori* understanding of how the model behaved with different values of reinforcements, we ran simulations on ranges of positive (1, 1.2, 1.4, 1.6, 1.8, 2) and negative (0, 0.1, 0.2, 0.3, 0.4, 0.5, 0.6, 0.7, 0.8, 0.9, 1) reinforcements for a total of 66 sets of parameters, for each environment type (one, two and three patches; Fig. S2). We simulated 10 environments for each environment type and computed 100 simulations of 50 foraging bouts per iteration (i.e. 1000 simulations per environment type and set of parameters).

Since our study focused on resource partitioning, we extracted the *Q_norm_* index values for these simulations and compared them. We plotted a heatmap showing the value of the mean *Q_norm_* index at the last foraging bout for each set of parameters and environment type (Fig. S3). In all types of environments, positive reinforcement had a strong effect on the final *Q_norm_* index value. High values of resource partitioning were obtained for positive reinforcement values > 1.5. By contrast, negative reinforcement only had an impact in environments with two or three patches. High values of resource partitioning were obtained for negative reinforcement values larger than 0.75.

For each pair of bees, we also looked at how this same index evolved over successive foraging bouts. Fig. S4 shows the dynamics of mean *Q_norm_* index across 50 foraging bouts for each combination of positive reinforcement (1.0, 1.2, 1.4, 1.6, 1.8, 2.0) and negative reinforcement (0, 0.1, 0.2, 0.3, 0.4, 0.5, 0.6, 0.7, 0.8, 0.9, 1) parameters, and each environment type (one patch, two patches, three patches). Higher values of both positive and negative reinforcements most often lead to faster resource partitioning (with some uncertainty due to the probabilistic nature of the model). Combinations of values in which the negative reinforcement factor was missing (violet gradient curves) led to a decrease in partitioning.

## Appendix S3. Qualitative comparison between simulations and observations in the narrow pentagon

We ran simulations with one forager to compare model outcomes to observational data using a second reference field study (Woodgate *et al*. 2017). In this study, the authors used five artificial flowers arranged in a narrow pentagon (Fig. S1B). Four bumblebees were tested during 27 to 61 foraging bouts each (flower visitation sequences were kindly provided by Joe Woodgate). One of these bumblebees was tested over different days and was therefore removed from the analyses (as bee’s experience memory drops overnight (Lihoreau *et al*. 2010)). In these conditions, none of the bumblebees developed a stable trapline, although all significantly increased their foraging efficiency with time (e.g. reduced travel distance and duration, increased similarity between two consecutive flower visitation sequences).

The sole implementation of positive reinforcement in Model 1 was sufficient to replicate the observations. While the use of the negative reinforcement alone in model 2 showed drastically different results, its addition with the positive reinforcement in model 3 had no major effect on route quality nor on route similarity trends (Fig. S5). Overall, simulations of models 1 and 3 showed good qualitative fit to the traplining behaviour observed in real bees – i.e. there is a trend of increasing route similarities across foraging bouts. Note however that the models tend to overestimate the bee ability to develop stable routes. This imperfect match could be due to the low amount of available experimental data in the original study (three individuals in Woodgate *et al*. 2017). Alternately, the model does not integrate any kind of stochastic exploration so that at each new step, bees do not provide the possibility to choose unknown spatial targets. Real bees, on the contrary, show phases of stochastic explorations during and after trapline formation (Woodgate *et al*. 2017; Kembro *et al*. 2019). Future experiments with more bees in a larger diversity of arrays of flowers will be useful to further quantitatively calibrate the model.

## Appendix S4. Predictions with more than two bees

We explored the emergence of resource partitioning in groups of 5 bees, and how this varies in environments containing 20, 25, 30, 40, 50, 70 and 100 flowers, thus encompassing a gradient of competition pressures from conditions where there are not enough flowers for all bees (20) to conditions where there are four times more flowers than necessary for all bees (100). For simplicity, flowers were evenly distributed (i.e. environment with one patch). The positive reinforcement factor was set at 1.5, and the negative reinforcement factor was set at 0.75. For each flower density, we generated 10 environments, and ran 100 simulations of 100 foraging bouts, for a total of 1000 simulations per density value. We computed the resource partitioning index (*Q_norm_*) at each foraging bout. The mean final *Q_norm_* was higher in environments with most flowers (Fig. S6). Plotting the mean final *Q_norm_* (final foraging bout) as a function of the number of available flowers confirmed that bees converge to a plateau where increasing the number of flowers only provokes an increase of partitioning (i.e. around 50 flowers) (Fig. S6).

## Appendix S5. Supplementary information on the similarity index

We used a route similarity index between two consecutive visitation sequences in order to assess how similar they were. We designed an index that would look both into (i) the similarity of vector uses throughout the bout, but also (ii) the similarity in sequence length.

This index works as follows. First it makes a census of all the vectors used in both sequences that will be called *a* and *b*. For the computation of this index, the “visits” of the nest are removed. Vectors used in both sequences *a* and *b* are determined. The flower visits that are in repeated vectors are then counted and summed as *s_ab_*. This sum is then divided by the number of visits in the longest sequence, hereafter *l_ab_*, multiplied by two. This multiplication by 2 in the denominator allows for accounting of length differences between the two sequences compared. The following equation is thus obtained:

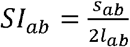

Example:

Sequence *a*: N 5 3 4 N
Sequence *b*: N 5 3 4 2 5 3 4 N

First, the visits to the nest are removed, giving the following sequences:

Sequence *a*: 5 3 4
Sequence *b*: 5 3 4 2 5 3 4

Then, the use of each vector in both sequences is quantified:

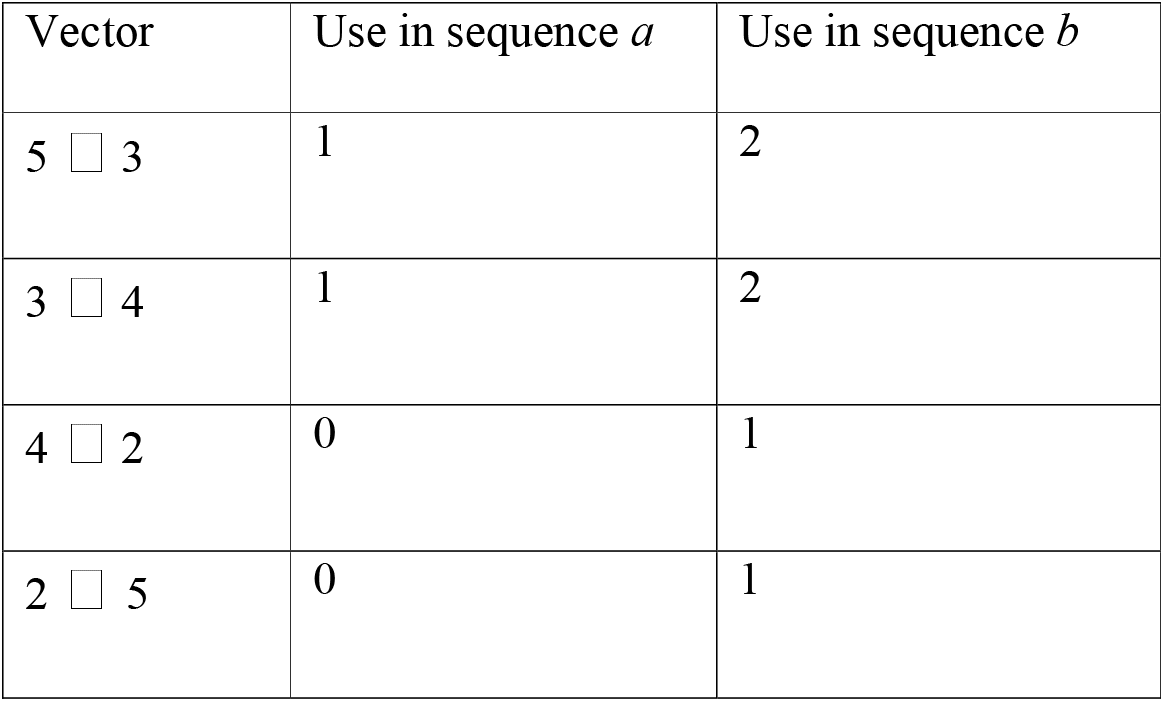

The vectors present in both sequences are identified (here in bold):

Sequence *a*: **5 3 4**
Sequence *b*: **5 3 4** 2 **5 3 4**

In this case, 9 total visits are part of repeated vectors. The longest sequence has a length of 7 visits.

Thus, the computation of our index is:

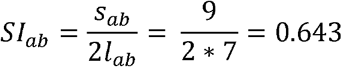

## Supplementary Figures

**Figure S1.**
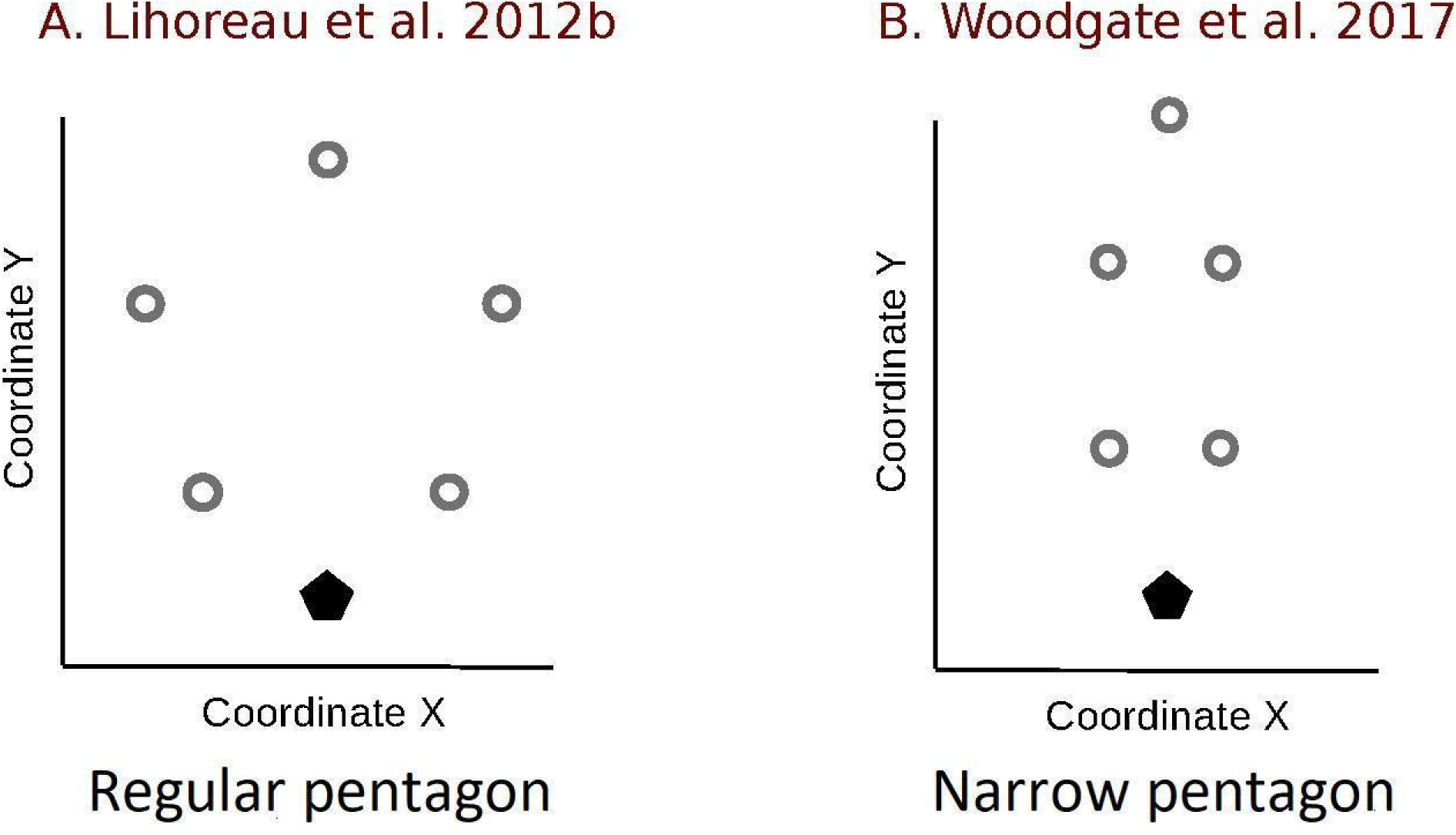
Arrays of artificial flowers (grey circles) and the colony nest (black pentagons) used to obtained the experimental datasets. A. Regular pentagon, modified from Lihoreau *et al*. (2012b). B. Narrow pentagon, modified from Woodgate *et al*. (2017).

**Figure S2.**
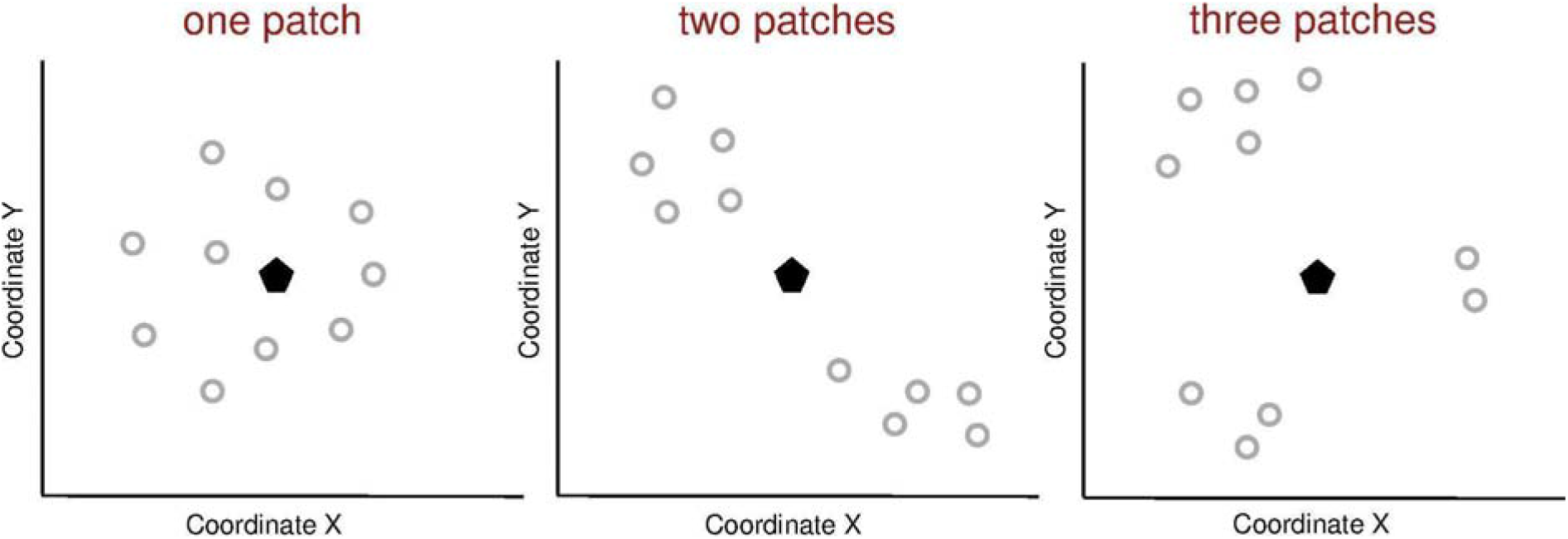
Examples of simulated environments. Spatial distribution of 10 flowers (grey circles) and a colony nest (black pentagon) in three types of environments defined by different levels of flower patchiness. A flower patch was characterized by: 1) a uniform distribution of flowers, 2) a lower distance between flowers within the patch than between all flowers from different patches (see details in methods).

**Figure S3.**
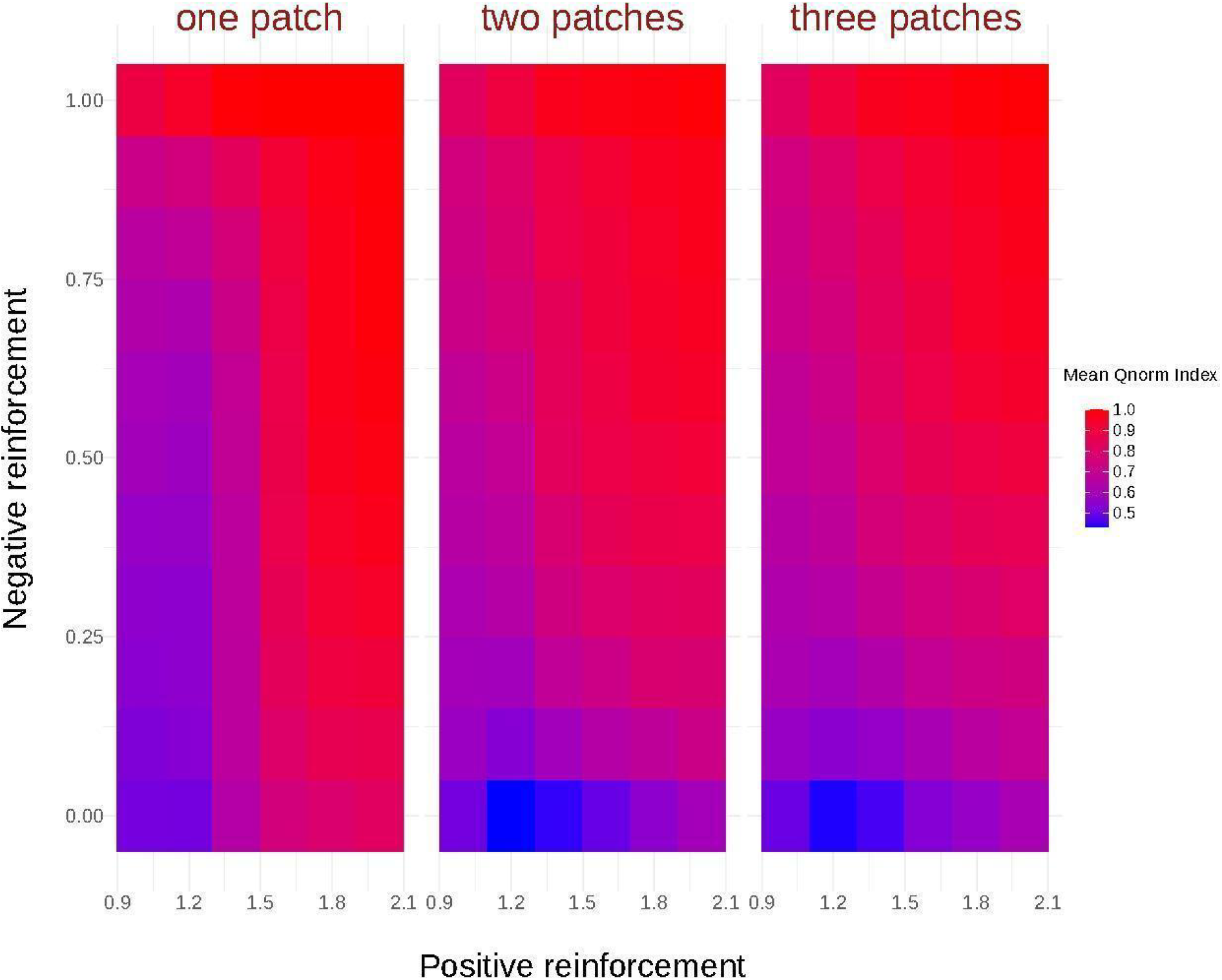
Heatmap showing the mean *Q_norm_* Index value after 50 foraging bouts (mean over 1000 simulations on 10 arrays of the same environment type), for each combination of positive reinforcement (1.0, 1.2, 1.4, 1.6, 1.8, 2.0) and negative reinforcement (0, 0.1, 0.2, 0.3, 0.4, 0.5, 0.6, 0.7, 0.8, 0.9, 1) parameters, and for each environment type (one patch, two patches, three patches). For simplicity, we inverted the values of negative reinforcement. 0 indicate models without negative reinforcement.

**Figure S4.**
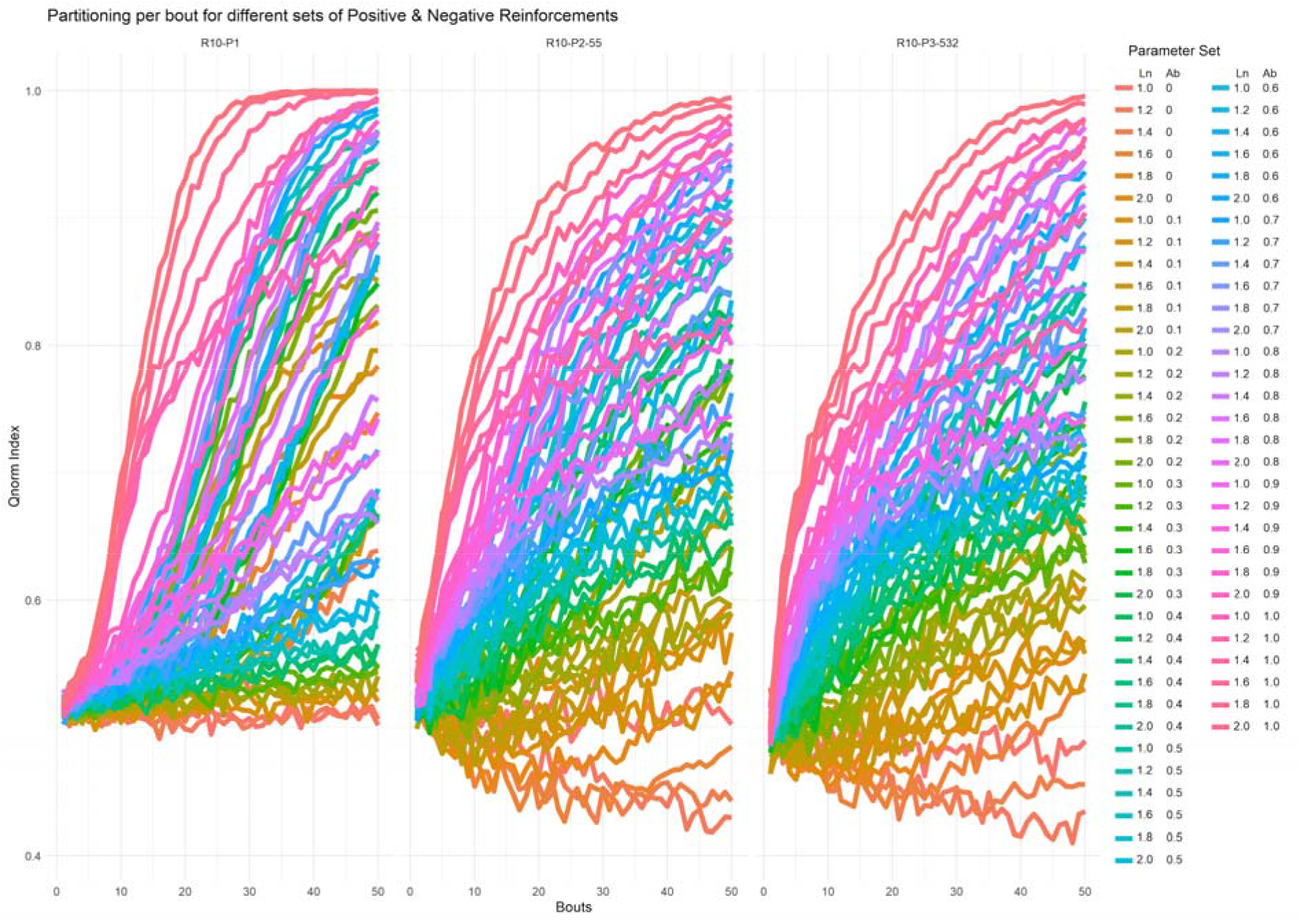
Dynamic of the mean *Q_norm_* Index across foraging bouts for each combination of positive (1.0, 1.2, 1.4, 1.6, 1.8, 2.0) and negative (0, 0.1, 0.2, 0.3, 0.4, 0.5, 0.6, 0.7, 0.8, 0.9, 1) reinforcement factors and for each environment type (one patch, two patches, three patches). For simplicity, we inverted the values of negative reinforcement here. 0 being models without negative reinforcement.

**Figure S5.**
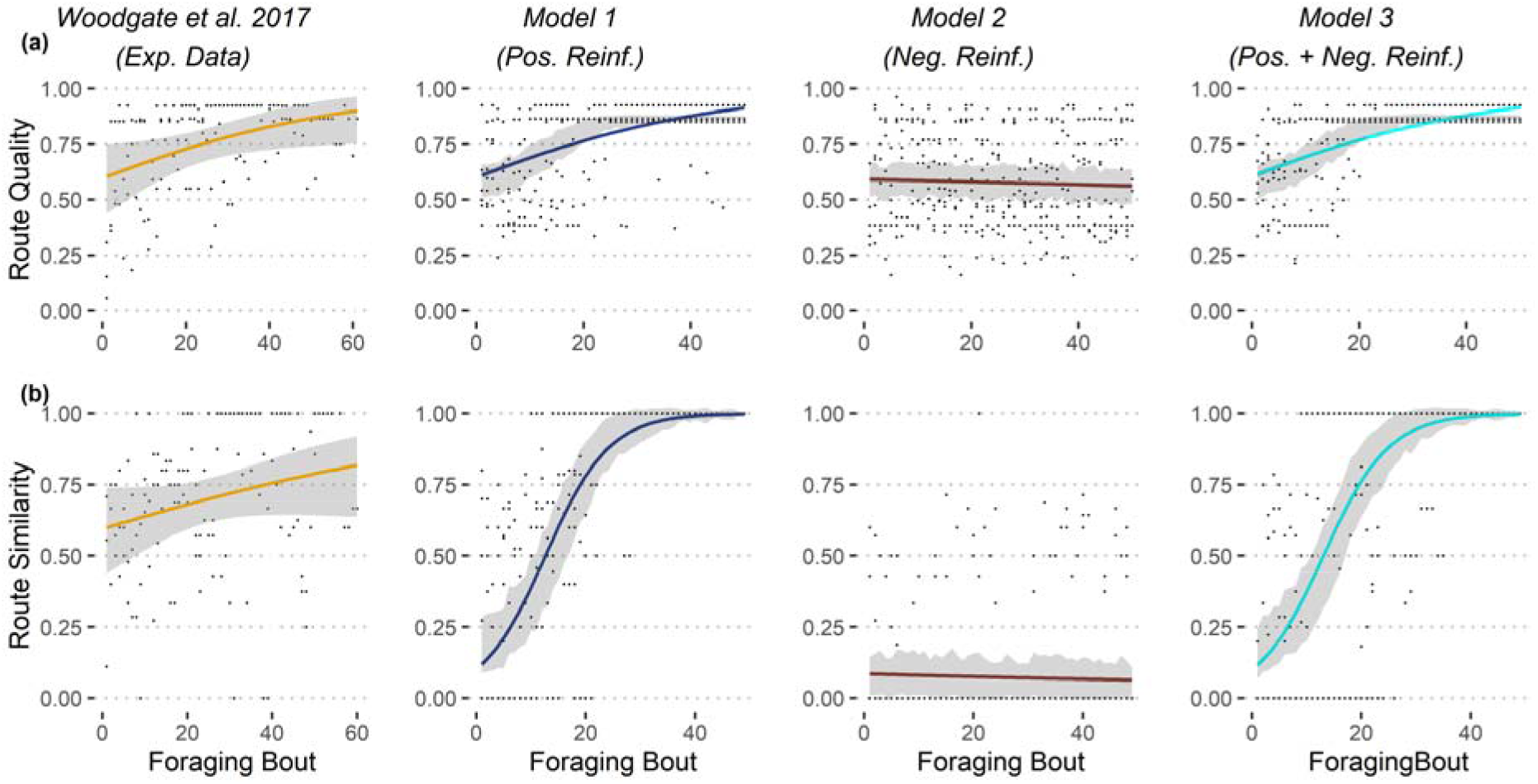
Qualitative comparisons of route qualities (A) and similarities (B) between simulations and experimental data (narrow pentagon array of flowers as in Woodgate *et al*. 2017) for one forager (see details of the models in Fig. 1). For each dataset, we show the estimated average trends across foraging bouts (colored curves), along with their 95% CI (gray areas). For simplicity, in the simulation plot we only represent a subsample of three random bees with their estimated 95% CI. Average trends were estimated using GLMM Binomial model with bee identity as random effect (bee identity nested in simulation identity for simulated data).

**Figure S6.**
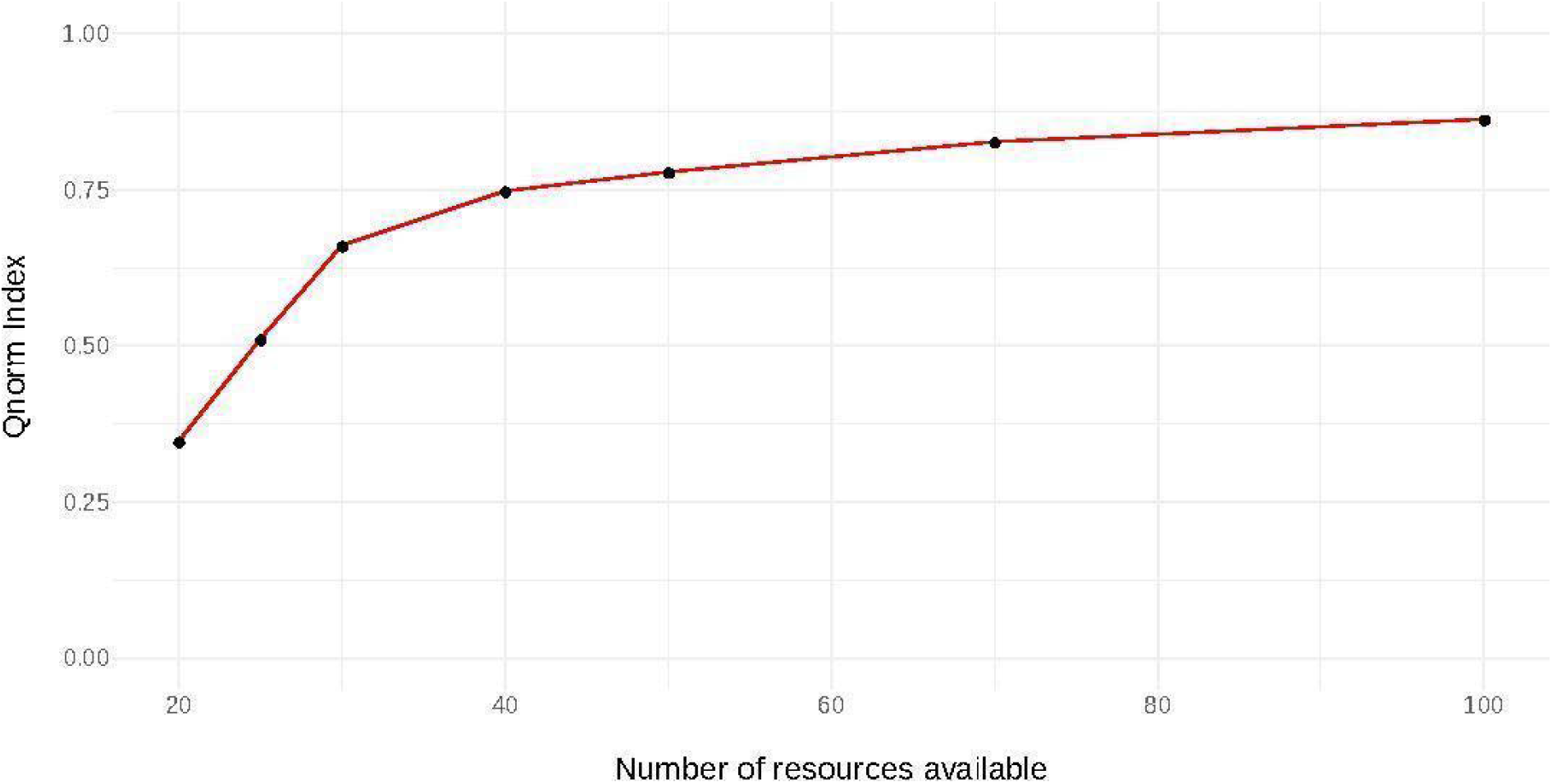
Evaluation of the mean final *Q_norm_* index (after 100 foraging bouts) as a function of increase resources availability. The model run has the positive reinforcement factor set at 1.5, and the negative reinforcement factor set at 0.75 with five bees foraging in environments of one patch.

**Video S1.** Example of simulation of five bees foraging in an environment with one patch of 50 flowers. Both positive and negative reinforcement rules are implemented (model 3). Bees performed 100 foraging bouts.

## References

Avarguès-Weber, A., de Brito Sanchez, M. G., Giurfa, M., & Dyer, A. G. 2010. Aversive reinforcement improves visual discrimination learning in free-flying honeybees. PloS One, 5(10). https://doi.org/10.1371/journal.pone.0015370

Bates, D., Mächler, M., Bolker, B., & Walker, S. 2015. Fitting linear mixed-effects models using lme4. Journal of Statistical Software, 67(1), 1–48. https://doi.org/10.18637/jss.v067.i01

Becher, M.A., Grimm, V., Thorbek, P., Horn, J., Kennedy, P.J., & Osborne, J.L. 2014. BEEHAVE: a systems model of honeybee colony dynamics and foraging to explore multifactorial causes of colony failure. Journal of Applied Ecology, 51(2), 470–482. https://doi.org/10.1111/1365-2664.12222

Becher, M.A., Grimm, V., Knapp, J., Horn, J., Twiston-Davies, G., & Osborne, J.L. 2016. BEESCOUT: A model of bee scouting behaviour and a software tool for characterizing nectar/pollen landscapes for BEEHAVE. Ecological Modelling, 24(11), 126–133. https://doi.org/10.1016/j.ecolmodel.2016.09.013

Becher, M.A., Twiston Davies, G., Penny, T.D., Goulson, D., Rotheray, E.L., & Osborne, J.L. 2018. Bumble BEEHAVE: A systems model for exploring multifactorial causes of bumblebee decline at individual, colony, population and community level. Journal of Applied Ecology, 55(6), 2790–2801. https://doi.org/10.1111/1365-2664.13165

Beckett, S.J. 2016. Improved community detection in weighted bipartite networks. Royal Society Open Science, 3(1). https://doi.org/10.1098/rsos.140536

Beshers, S. N., & Fewell, J. H. 2001. Models of division of labor in social insects. Annual Review of Entomology, 46(1), 413–440. https://doi.org/10.1146/annurev.ento.46.1.413

Buatois, A., & Lihoreau, M. 2016. Evidence of trapline foraging in honeybees. Journal of Experimental Biology, 219(16), 2426–2429. https://doi.org/10.1242/jeb.143214

Chittka, L., Dyer, A.G., Bock, F., & Dornhaus, A. 2003. Bees trade off foraging speed for accuracy. Nature, 424(6947), 388–388. https://doi.org/10.1038/424388a

Dawson, E. H., & Chittka, L. 2012. Conspecific and heterospecific information use in bumblebees. PloS One, 7(2). https://doi.org/10.1371/journal.pone.0031444

Dormann, C.F., Gruber, B., & Fründ, J. 2008. Introducing the bipartite package: analysing ecological networks. R News, 8(2), 8–11.

Dormann C.F., & Strauss R. 2014. A method for detecting modules in quantitative bipartite networks. Methods in Ecology and Evolution, 5(1), 90–98. https://doi.org/10.1111/2041-210X.12139

Dunlap, A.S., Nielsen, M.E., Dornhaus, A., & Papaj, D.R. 2016. Foraging bumble bees weigh the reliability of personal and social information. Current Biology, 26(9), 1195–1199. https://doi.org/10.1016/j.cub.2016.03.009

Fretwell, S.D., & Lucas, H.L. 1969. On territorial behavior and other factors influencing habitat distribution in birds. Acta Biotheoretica, 19(1), 16–36. https://doi.org/10.1007/BF01601955

Fründ, J., Linsenmair, K.E., & Blüthgen, N. 2010. Pollinator diversity and specialization in relation to flower diversity. Oikos, 119(10), 1581–1590. https://doi.org/10.1111/j.1600-0706.2010.18450.x

Fründ, J., Dormann, C.F., Holzschuh, A., & Tscharntke, T. 2013. Bee diversity effects on pollination depend on functional complementarity and niche shifts. Ecology, 94(9), 2042–2054. https://doi.org/10.1890/12-1620.1

Garrison, L. K., Kleineidam, C. J., & Weidenmüller, A. 2018. Behavioral flexibility promotes collective consistency in a social insect. Scientific Reports, 8(1), 1–11. https://doi.org/10.1038/s41598-018-33917-7

Giraldeau, L.A., & Caraco, T. 2000. Social Foraging Theory, Princeton University Press, Princeton.

Giurfa, M. 2004. Conditioning procedure and color discrimination in the honeybee *Apis mellifera*. Naturwissenschaften, 91(5), 228–231.

Giurfa, M., 2013. Cognition with few neurons: higher-order learning in insects. Trends in neurosciences, 36(5), 285–294. https://doi.org/10.1016/j.tins.2012.12.011

Goldshtein, A., Handel, M., Eitan, O., Bonstein, A., Shaler, T., Collet, S., Greif, S., Medellin, R. A., Emek, Y., Korman, A., & Yovel, Y. 2020. Reinforcement learning enables resource partitioning in foraging bats. Current Biology, 30(20), 4096–4102. https://doi.org/10.1016/j.cub.2020.07.079

Goulson, D. 2010. Bumblebees: Behaviour, Ecology, and Conservation. Oxford University Press, Oxford.

Grimm, V., Berger, U., Bastiansen, F., Eliassen, S., Ginot, V., Giske, J., Goss-Custard, J., Grand, T., Heinz, S. K., Huse, G., Huth, A., Jepsen, J. U., Jørgensen, C., Mooij, W. M., Müller, B., Pe’er, G., Piou, C., Railsback, S. F., Robbins, A. M., Robbins, M. M., Rossmanith, E., Rüger, N., Strand, E., Souissi, S., Stillman, R. A., Vabø, R., Visser, U., & DeAngelis, D. L. 2006. A standard protocol for describing individual-based and agent-based models. Ecological Modelling, 198(1-2), 115–126. https://doi.org/10.1016/j.ecolmodel.2006.04.023

Grimm, V., Railsback, S. F., Vincenot, C. E., Berger, U., Gallagher, C., DeAngelis, D. L., Edmonds, B., Ge, J., Giske, J., Groeneveld, J., Johnston, A. S. A., Milles, A., Nabe-Nielsen, J., Polhill, J. G., Radchuk, V., Rohwäder, M. S., Stillman, R. A., Thiele, J. C., & Ayllón, D. 2020. The ODD protocol for describing agent-based and other simulation models: A second update to improve clarity, replication, and structural realism. Journal of Artificial Societies and Social Simulation, 23(2). https://doi.org/10.18564/jasss.4259

Hendriksma, H.P., Toth, A.L., & Shafir, S. 2019. Individual and colony level foraging decisions of bumble bees and honey bees in relation to balancing of nutrient needs. Frontiers in Ecology and Evolution, 7(1), 177. https://doi.org/10.3389/fevo.2019.00177

Janzen, D.H. 1971. Euglossine bees as long-distance pollinators of tropical plants. Science, 171(3967), 203–205.

Johst, K., Berryman, A., & Lima, M. 2008. From individual interactions to population dynamics: individual resource partitioning simulation exposes the causes of nonlinear intra-specific competition. Population Ecology, 50(1), 79–90. https://doi.org/10.1007/s10144-007-0061-5

Kazakova, V. A., Wu, A. S., & Sukthankar, G. R. 2020. Respecializing swarms by forgetting reinforced thresholds. Swarm Intelligence 14(3), 171–204. https://doi.org/10.1007/s11721-020-00181-3

Kembro, J.M., Lihoreau, M., Garriga, J., Raposo, E.P., & Bartumeus, F. 2019. Bumblebees learn foraging routes through exploitation–exploration cycles. Journal of the Royal Society Interface, 16(156). https://doi.org/10.1098/rsif.2019.0103

Klein, S., Pasquaretta, C., Barron, A.B., Devaud, J.M., & Lihoreau, M. 2017. Inter-individual variability in the foraging behaviour of traplining bumblebees. Scientific Reports, 7(1), 1–12. https://doi.org/10.1038/s41598-017-04919-8

Kraus, S., Gómez-Moracho, T., Pasquaretta, C., Latil, G., Dussutour, A., & Lihoreau, M. 2019. Bumblebees adjust protein and lipid collection rules to the presence of brood. Current Zoology, 65(4), 437–446. https://doi.org/10.1093/cz/zoz026

Le Moël, F., Stone, T.J., Lihoreau, M., Wystrach, A., & Webb, B. 2019. The central complex as a potential substrate for vector-based navigation. Frontiers in Psychology 10(1). https://doi.org/10.3389/fpsyg.2019.00690

Leadbeater, E., & Chittka, L. 2005. A new mode of information transfer in foraging bumblebees? Current Biology, 15(12), 447–448.

Leadbeater, E., & Chittka, L. 2011. Do inexperienced bumblebee foragers use scent marks as social information? Animal Cognition 14(6), 915–919. https://doi.org/10.1007/s10071-011-0423-4

Lenth, R., Singmann, H., Love, J., Buerkner, P., & Herve, M. 2019. Estimated marginal means, aka least-squares means. R package version 1.3. 2.

Lihoreau, M., Chittka, L., & Raine, N.E. 2010. Travel optimization by foraging bumblebees through readjustments of traplines after discovery of new feeding locations. The American Naturalist 176(6), 744–757. https://doi.org/10.1086/657042

Lihoreau, M., Chittka, L., & Raine, N. E. 2016. Monitoring flower visitation networks and interactions between pairs of bumble bees in a large outdoor flight cage. PloS One, 11(3). https://doi.org/10.1371/journal.pone.0150844

Lihoreau, M., Chittka, L., Le Comber, S.C., & Raine, N.E. 2012a. Bees do not use nearest-neighbour rules for optimization of multi-location routes. Biology Letters, 8(1), 13–16. https://doi.org/10.1098/rsbl.2011.0661

Lihoreau, M., Raine, N.E., Reynolds, A.M., Stelzer, R.J., Lim, K.S., Smith, A.D., Osborne, J. L., & Chittka, L. 2012b. Radar tracking and motion-sensitive cameras on flowers reveal the development of pollinator multi-destination routes over large spatial scales. PloS Biology, 10(9). https://doi.org/10.1371/journal.pbio.1001392

Makino, T.T., & Sakai, S. 2005. Does interaction between bumblebees *(Bombus ignitus)* reduce their foraging area?: Bee-removal experiments in a net cage. Behavioural Ecology and Sociobiology, 57(6), 617–622. https://doi.org/10.1007/s00265-004-0877-3

Makino, T.T. 2013. Longer visits on familiar plants?: testing a regular visitor’s tendency to probe more flowers than occasional visitors. Naturwissenschaften, 100(7), 659–666. https://doi.org/10.1007/s00114-013-1062-1

Menzel, R. 1990. Learning, memory, and “cognition” in honey bees. Neurobiology of Comparative Cognition. Ed. Kesner, R.P. and Olton, D.S. Lawrence Erlbaum Associates, Inc., Publishers. 237–292.

Nagamitsu, T., & Inoue, T. 1997. Aggressive foraging of social bees as a mechanism of floral resource partitioning in an Asian tropical rainforest. Oecologia, 110(3), 432–439.

Ohashi, K., & Thomson, J. D. 2005. Efficient harvesting of renewing resources. Behavioral Ecology, 16(3), 592–605. https://doi.org/10.1093/beheco/ari031

Ohashi, K., Thomson, J. D., & D’souza, D. 2007. Trapline foraging by bumble bees: IV. Optimization of route geometry in the absence of competition. Behavioral Ecology, 18(1), 1–11. https://doi.org/10.1093/beheco/arl053

Ohashi, K., Leslie, A., & Thomson, J.D. 2008. Trapline foraging by bumble bees: V. Effects of experience and priority on competitive performance. Behavioral Ecology, 19(5), 936–948. https://doi.org/10.1093/beheco/arn048

Ohashi, K., & Thomson, J. D. 2009. Trapline foraging by pollinators: its ontogeny, economics and possible consequences for plants. Annals of Botany, 103(9), 1365–1378. https://doi.org/10.1093/aob/mcp088

Osborne, J.L., Clark, S.J., Morris, R.J., Williams, I.H., Riley, J.R., Smith, A.D., Reynolds, D. R., & Edwards, A.S. 1999. A landscape scale study of bumble bee foraging range and constancy, using harmonic radar. Journal of Applied Ecology, 36(4), 519–533. https://doi.org/10.1046/j.1365-2664.1999.00428.x

Pasquaretta, C., Jeanson, R., Andalo, C., Chittka, L., & Lihoreau, M., 2017. Analysing plant-pollinator interactions with spatial movement networks. Ecological Entomology. 42(Suppl. 1), 4–17. https://doi.org/10.1111/een.12446

Pasquaretta, C., & Jeanson, R. 2018. Division of labor as a bipartite network. Behavioral Ecology, 29(2), 342–352. https://doi.org/10.1093/beheco/arx170

Pasquaretta, C., Jeanson, R., Pansanel, J., Raine, N.E., Chittka, L., & Lihoreau, M. 2019. A spatial network analysis of resource partitioning between bumblebees foraging on artificial flowers in a flight cage. Movement Ecology, 7(1), 1–10. https://doi.org/10.1186/s40462-019-0150-z

Possingham, H. P. 1989. The distribution and abundance of resources encountered by a forager. The American Naturalist, 133(1), 42–60. https://doi.org/10.1086/284900

R Core Team 2018. R: A language and environment for statistical computing. R Foundation for Statistical Computing, Vienna, Austria. URL https://www.R-project.org/.

Raine, N.E., & Chittka, L. 2012. No trade-off between learning speed and associative flexibility in bumblebees: a reversal learning test with multiple colonies. PloS One, 7(9). https://doi.org/10.1371/journal.pone.0045096

Reynolds, A.M., Lihoreau, M., & Chittka, L. 2013. A simple iterative model accurately captures complex trapline formation by bumblebees across spatial scales and flower arrangements. PLoS Computational Biology, 9(3). https://doi.org/10.1371/journal.pcbi.1002938

Robinson, E. J., Jackson, D. E., Holcombe, M., & Ratnieks, F. L. 2005. ‘No entry’ signal in ant foraging. Nature, 438(7067), 442–442. https://doi.org/10.1038/438442a

Saleh, N., & Chittka, L. 2007. Traplining in bumblebees *(Bombus impatiens):* a foraging strategy’s ontogeny and the importance of spatial reference memory in short-range foraging. Oecologia, 151(4), 719–730. https://doi.org/10.1007/s00442-006-0607-9

Schwaerzel, M., Monastirioti, M., Scholz, H., Friggi-Grelin, F., Birman, S., & Heisenberg, M. 2003. Dopamine and octopamine differentiate between aversive and appetitive olfactory memories in Drosophila. Journal of Neuroscience, 23(33), 10495–10502. https://doi.org/10.1523/JNEUROSCI.23-33-10495.2003

Seeley, T. D., Visscher, P. K., Schlegel, T., Hogan, P. M., Franks, N. R., & Marshall, J. A. 2012. Stop signals provide cross inhibition in collective decision-making by honeybee swarms. Science, 335(6064), 108–111. https://doi.org/10.1126/science.1210361

Stone, T., Webb, B., Adden, A., Weddig, N.B., Honkanen, A., Templin, R., Wcislo, W., Scimeca, L., Warrant, E., & Heinze, S. 2017. An anatomically constrained model for path integration in the bee brain. Current Biology, 27(20), 3069–3085. https://doi.org/10.1016/j.cub.2017.08.052

Stout, J. C., & Goulson, D. 2001. The use of conspecific and interspecific scent marks by foraging bumblebees and honeybees. Animal Behaviour, 62(1), 183–189. https://doi.org/10.1006/anbe.2001.1729

Sumpter, D. J. 2010. Collective animal Animal Behavior. Princeton University Press.

Thomson, J.D. 1996. Trapline foraging by bumblebees: I. Persistence of flight-path geometry. Behavioral Ecology, 7(2), 158–164. https://doi.org/10.1093/beheco/7.2.158

Thomson, J.D., Slatkin, M., & Thomson, B.A. 1997. Trapline foraging by bumble bees: II. Definition and detection from sequence data. Behavioral Ecology 8(2), 199–210. https://doi.org/10.1093/beheco/8.2.199

Tinker, T.M., Guimaraes Jr, P.R., Novak, M., Marquitti, F.M.D., Bodkin, J.L., Staedler, M., Bentall, M., & Estes, J.A. 2012. Structure and mechanism of diet specialization: testing models of individual variation in resource use with sea otters. Ecology Letters, 15(5), 475–483. https://doi.org/10.1111/j.1461-0248.2012.01760.x

Valdovinos, F. S., Brosi, B. J., Briggs, H. M., Moisset de Espanés, P., Ramos Jiliberto, R., & Martinez, N. D. 2016. Niche partitioning due to adaptive foraging reverses effects of nestedness and connectance on pollination network stability. Ecology Letters, 19(10), 1277–1286. https://doi.org/10.1111/ele.12664

Vergoz, V., Roussel, E., Sandoz, J.C., & Giurfa, M. 2007. Aversive learning in honeybees revealed by the olfactory conditioning of the sting extension reflex. PloS One, 2(3). https://doi.org/10.1371/journal.pone.0000288

Wolf, S., & Moritz, R.F. 2008. Foraging distance in *Bombus terrestris* L. (Hymenoptera: Apidae). Apidologie, 39(4), 419–427. https://doi.org/10.1051/apido:2008020

Woodgate, J. L., Makinson, J. C., Lim, K. S., Reynolds, A. M., & Chittka, L. 2016. Life-long radar tracking of bumblebees. PloS One, 11(8). https://doi.org/10.1371/journal.pone.0160333

Woodgate, J.L., Makinson, J.C., Lim, K.S., Reynolds, A.M., & Chittka, L. 2017. Continuous radar tracking illustrates the development of multi-destination routes of bumblebees. Scientific Reports, 7(1), 1–15. https://doi.org/10.1038/s41598-017-17553-1

Wright, G.A., Nicolson, S.W., & Shafir, S., 2018. Nutritional physiology and ecology of honey bees. Annual Review of Entomology, 63(1), 327–344. https://doi.org/10.1146/annurev-ento-020117-043423

Yokoi, T., & Fujisaki, K., 2009. Recognition of scent marks in solitary bees to avoid previously visited flowers. Ecological Research, 24(4), 803–809. https://doi.org/10.1007/s11284-008-0551-8

