## Supplementary materials for "A model of resource partitioning between foraging bees based on positive and negative associations"

**Appendix S1.** *ODD Protocol.*

Below we provide a description of our models following the ODD (Overview, Design concepts, Details) protocol (Grimm et al. 2006, 2020).

**1 – Purpose and patterns**

The purpose of the models is to offer an explanation to how multiple bees learn to optimize their foraging efficiency. Bees are expected to do so by developing efficient routes (traplines) minimizing spatial overlaps with other foragers (resource partitioning) (Lihoreau et al., 2016). We suggest such process can be achieved through combinations of positive and negative reinforcements during flower visits.

**2 - Entities, state variables and scales**

The models depict three kinds of entities: bees, flowers and the colony nest. Bees go out foraging for nectar rewards on flowers and return to the nest. Numbers of bees and flowers can be adjusted in the model. In our simulations we set a ratio of five flowers per bee as to fit the experimental data we replicated (Lihoreau et al., 2012; Woodgate et al., 2016). A single colony nest is represented, from which all bees forage and come back to. The bees are defined by the following set of state variables:

| Variable name | Variable type and units | Meaning |
| --- | --- | --- |
| *ID* | Integer, constant | A simple number to identify each bee in the model. |
| *crop* | Integer, dynamic | A count of how many flower rewards (volume of nectar) the bee has collected since its departure from the nest. |
| *maxCrop* | Integer, constant | A maximum number of rewards (volume of nectar) a bee can hold at a time. |
| *probabilityMatrix* | Matrix of probabilities, dynamic | A square matrix of length equal to the sum of number of flowers and the nest, depicting the probability to move between each entity. |
| *indPos* | Integer, dynamic | An integer showing the ID of the flower currently visited by the bee. |
| *winProbabilities* | Float, constant | Value indicative of the probability of winning an encounter on a flower (competition by interference). |

The flowers are static entities placed in the environment. They are defined by:

| Variable name | Variable type and units | Meaning |
| --- | --- | --- |
| *ID* | Integer, constant | A simple number to identify each flower in the model. |
| *resourceOnFlower* | Integer, dynamic | Variable showing the resource availability on the flower (0 or 1). |
| *x,y* | Floats, constant | Coordinates of the flower in the environment. |

The nest is a unique entity which is represented by the following variables:

| Variable name | Variable type and units | Meaning |
| --- | --- | --- |
| *x,y* | Floats, constant | Coordinates of the nest in the environment. |

Both the spatial and temporal scales are represented. Space is represented by the relative distance between the different flowers. Internal parameters define min/max ranges in which each entity can be placed, relative to other entities (see *CreateEnvironment* submodel). Time is represented abstractly. At each step, the bees visit a flower, and possibly feed on it. We assume all travel and flower manipulation times are identical.

*arrayGeometry*:

| ID | x | y |
| --- | --- | --- |
| 0 | 25 | -16.5831239518 |
| 1 | 0 | 0 |
| 2 | -15.4508497187 | 47.5528258148 |
| 3 | 25 | 76.9420884294 |
| 4 | 65.4508497187 | 47.5528258148 |
| 5 | 50 | 0 |

*arrayInfos*

| environmentType | numberOfResources | numberOfPatches | patchinessIndex | envSize | flowerPerPatch |
| --- | --- | --- | --- | --- | --- |
| lihoreau2012 | 5 | 1 | NA | NA | NULL |

| Vector | Use in sequence *a* | Use in sequence *b* |
| --- | --- | --- |
| 5 ➝ 3 | 1 | 2 |
| 3 ➝ 4 | 1 | 2 |
| 4 ➝ 2 | 0 | 1 |
| 2 ➝ 5 | 0 | 1 |

The vectors present in both sequences are identified (here in bold):

Sequence *a*: **5 3 4**

Sequence *b*: **5 3 4** 2 **5 3 4**

In this case, 9 total visits are part of repeated vectors. The longest sequence has a length of 7 visits. Thus, the computation of our index is:

$${SI}_{ab}=\frac{s_{ab}}{{2l}_{ab}}= \frac{9}{2*7}=0.643$$

**Supplementary Figures**

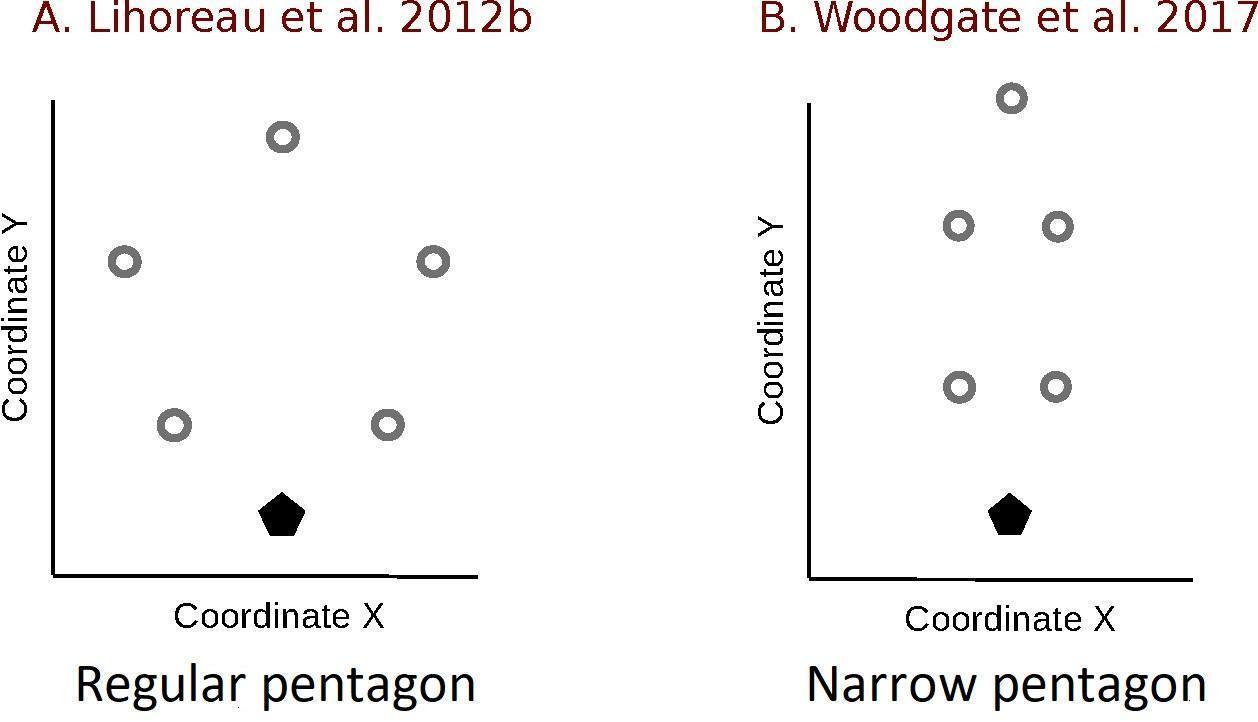

**Figure S1.** Arrays of artificial flowers (grey circles) and the colony nest (black pentagons) used to obtained the experimental datasets. A. Regular pentagon, modified from Lihoreau *et al.* (2012b). B. Narrow pentagon, modified from Woodgate *et al.* (2017).

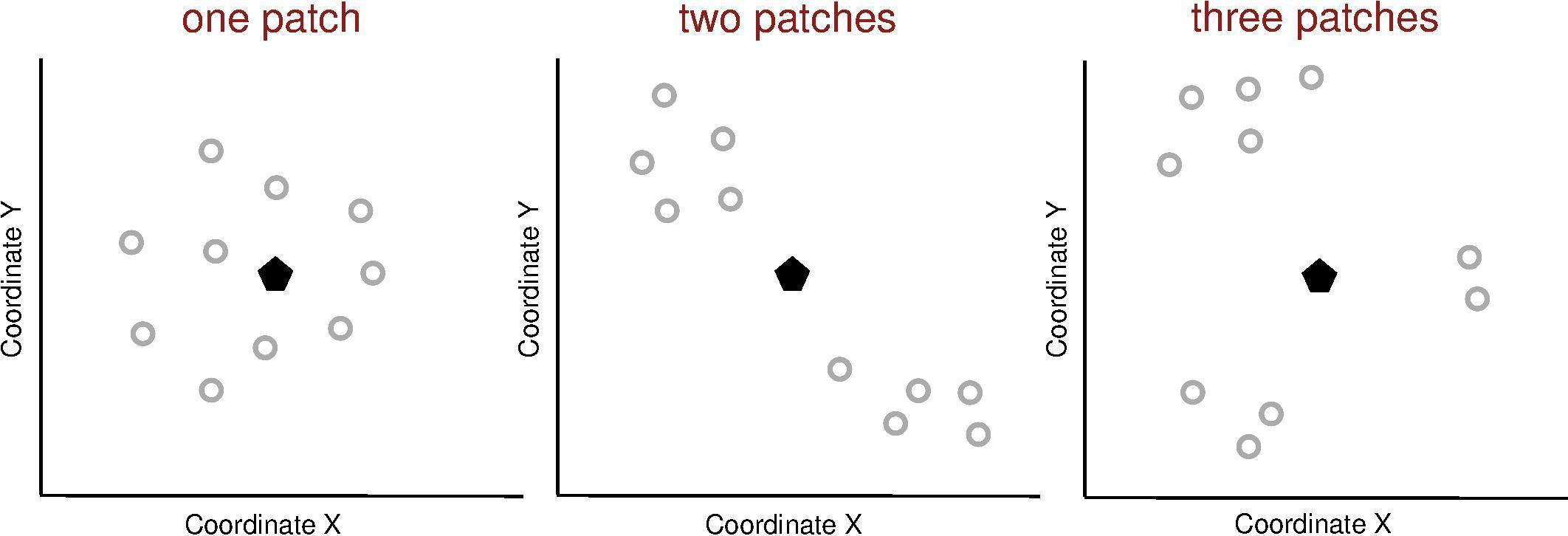

**Figure S2.** Examples of simulated environments. Spatial distribution of 10 flowers (grey circles) and a colony nest (black pentagon) in three types of environments defined by different levels of flower patchiness. A flower patch was characterized by: 1) a uniform distribution of flowers, 2) a lower distance between flowers within the patch than between all flowers from different patches (see details in methods).

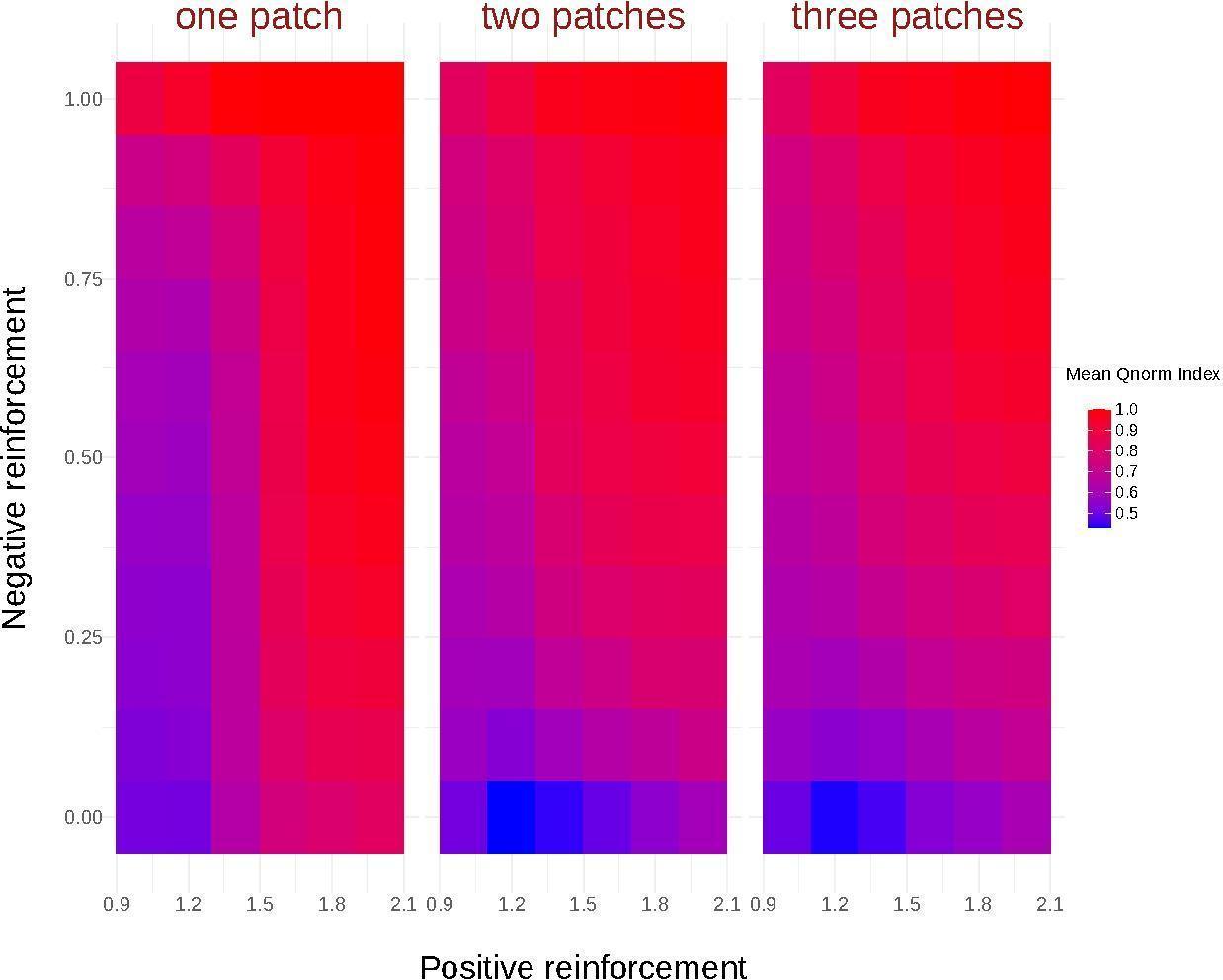

**Figure S3.** Heatmap showing the mean $Q_{norm}$ Index value after 50 foraging bouts (mean over 1000 simulations on 10 arrays of the same environment type), for each combination of positive reinforcement (1.0, 1.2, 1.4, 1.6, 1.8, 2.0) and negative reinforcement (0, 0.1, 0.2, 0.3, 0.4, 0.5, 0.6, 0.7, 0.8, 0.9, 1) parameters, and for each environment type (one patch, two patches, three patches). For simplicity, we inverted the values of negative reinforcement. 0 indicate models without negative reinforcement.

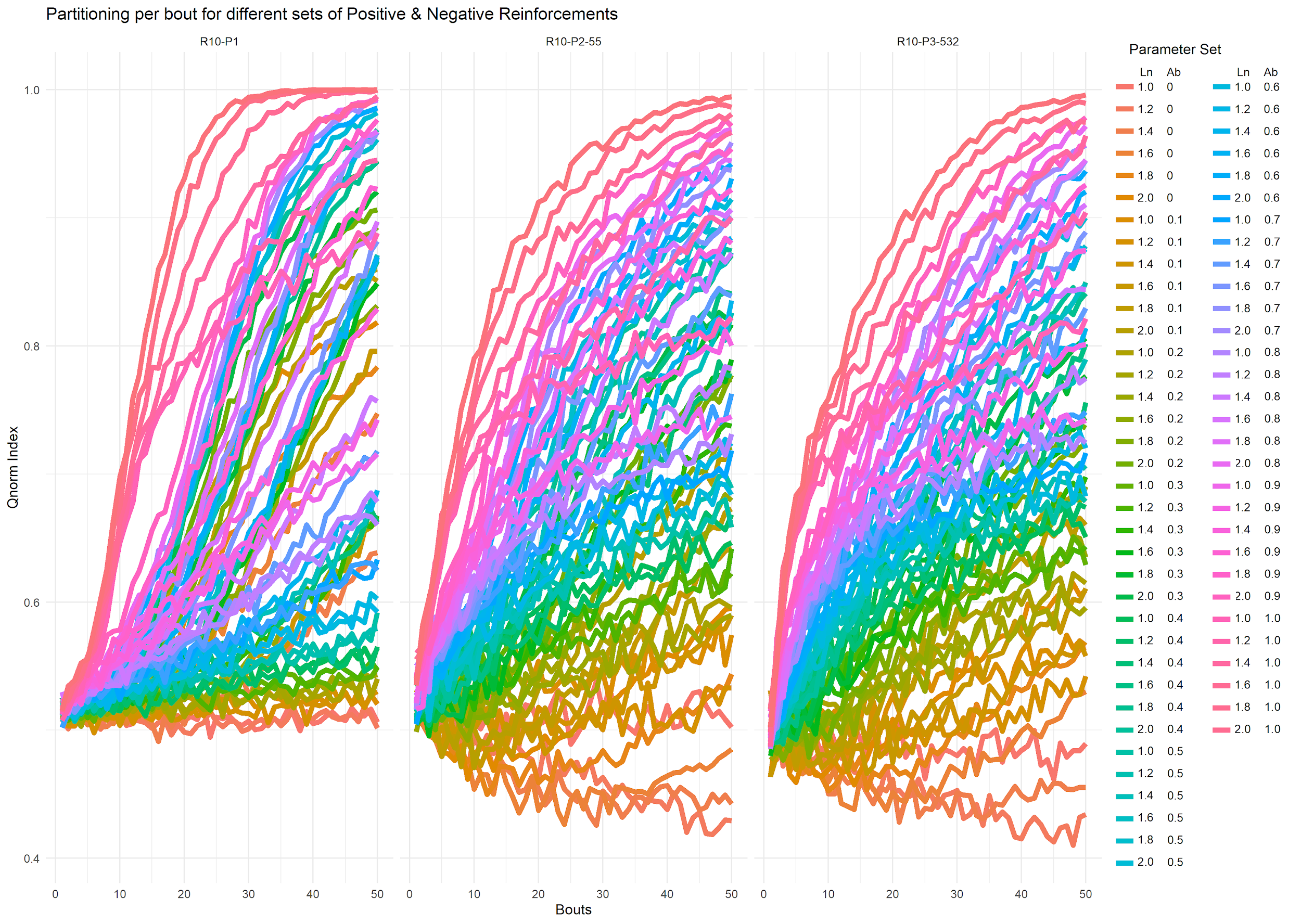

**Figure S4.** Dynamic of the mean $Q_{norm}$ Index across foraging bouts for each combination of positive (1.0, 1.2, 1.4, 1.6, 1.8, 2.0) and negative (0, 0.1, 0.2, 0.3, 0.4, 0.5, 0.6, 0.7, 0.8, 0.9, 1) reinforcement factors and for each environment type (one patch, two patches, three patches). For simplicity, we inverted the values of negative reinforcement here. 0 being models without negative reinforcement.

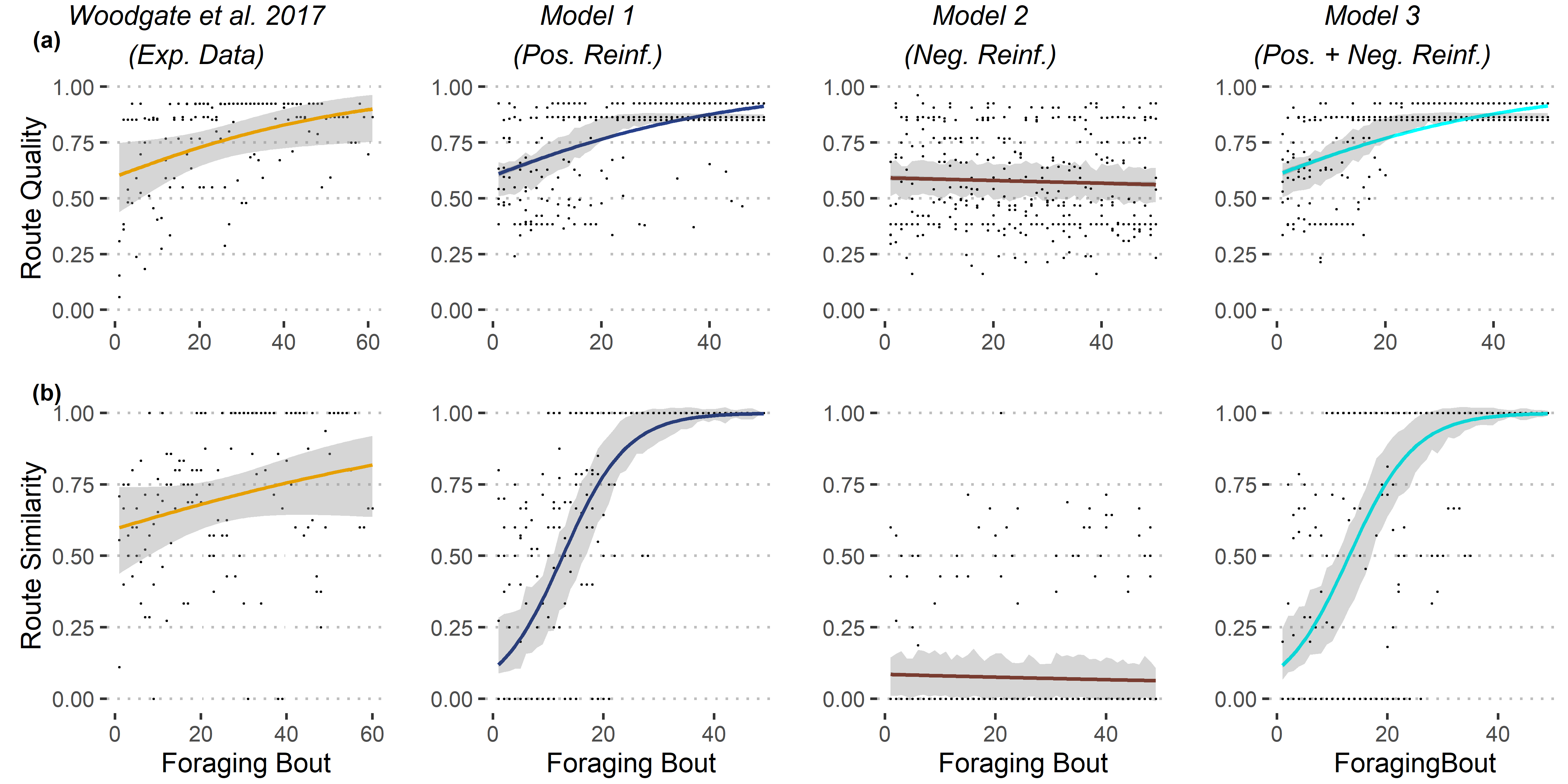

**Figure S5.** Qualitative comparisons of route qualities (A) and similarities (B) between simulations and experimental data (narrow pentagon array of flowers as in Woodgate *et al.* 2017) for one forager (see details of the models in Fig. 1). For each dataset, we show the estimated average trends across foraging bouts (colored curves), along with their 95% CI (gray areas). For simplicity, in the simulation plot we only represent a subsample of three random bees with their estimated 95% CI. Average trends were estimated using GLMM Binomial model with bee identity as random effect (bee identity nested in simulation identity for simulated data).

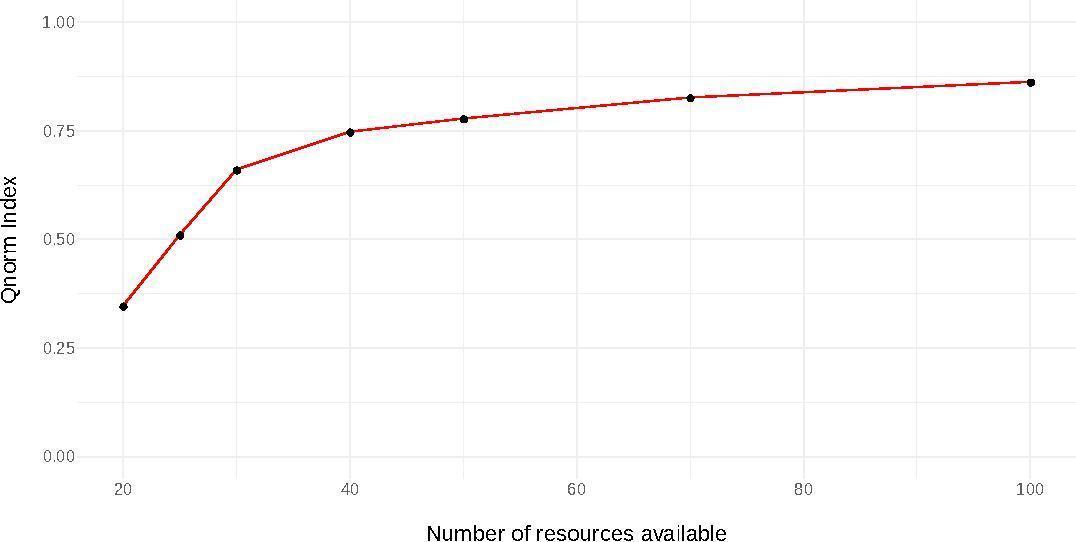

**Figure S6.** Evaluation of the mean final $Q_{norm}$ index (after 100 foraging bouts) as a function of increase resources availability. The model run has the positive reinforcement factor set at 1.5, and the negative reinforcement factor set at 0.75 with five bees foraging in environments of one patch.

**Video S1.** Example of simulation of five bees foraging in an environment with one patch of 50 flowers. Both positive and negative reinforcement rules are implemented (model 3). Bees performed 100 foraging bouts.
